# Direct delivery of stabilized Cas-embedded base editors achieves efficient and accurate editing of clinically relevant targets

**DOI:** 10.1101/2024.02.08.579528

**Authors:** Jeong Min Lee, Jing Zeng, Pengpeng Liu, My Anh Nguyen, Diego Suchenski Loustaunau, Daniel E. Bauer, Nese Kurt Yilmaz, Scot A. Wolfe, Celia A. Schiffer

**Author notes:** Joint last author (Nese Kurt Yilmaz, Scot A. Wolfe, Celia A. Schiffer).

## Abstract

Over the last 5 years, cytosine base editors (CBEs) have emerged as a promising therapeutic tool for specific editing of single nucleotide variants and disrupting specific genes associated with disease. Despite this promise, the currently available CBE’s have the significant liabilities of off-target and bystander editing activities, in part due to the mechanism by which they are delivered, causing limitations in their potential applications. In this study we engineered highly stabilized Cas-embedded CBEs (sCE_CBEs) that integrate several recent advances, and that are highly expressible and soluble for direct delivery into cells as ribonucleoprotein (RNP) complexes. Our resulting sCE_CBE RNP complexes efficiently and specifically target TC dinucleotides with minimal off-target or bystander mutations. Additional uracil glycosylase inhibitor (UGI) protein in *trans* further increased C-to-T editing efficiency and target purity in a dose-dependent manner, minimizing indel formation to untreated levels. A single electroporation was sufficient to effectively edit the therapeutically relevant locus for sickle cell disease in hematopoietic stem and progenitor cells (HSPC) in a dose dependent manner without cellular toxicity. Significantly, these sCE_CBE RNPs permitted for the transplantation of edited HSPCs confirming highly efficient editing in engrafting hematopoietic stem cells in mice. The success of the designed sCBE editors, with improved solubility and enhanced on-target editing, demonstrates promising agents for cytosine base editing at other disease-related sites in HSPCs and other cell types.

## Introduction

Cytosine base editors (CBEs) convert C•G base pairs to T•A base pairs within a small editing window specified by a guide RNA (sgRNA), allowing for single base changes in the genomic target (1–3). Compared to the CRISPR-Cas9 gene editing system which induces double-strand breaks (DSBs), CBEs are more efficient and carry less risk of genomic alterations such as insertions, deletions, and translocations (4). CBEs have, therefore, emerged as a popular tool for specific editing of single nucleotide variants (SNVs) and for disrupting specific genes associated with disease (5). However, despite recent progress and the large number of different versions of CBEs reported over the last 5 years (3, 6–9), the currently available CBEs still have the significant liabilities of off-target and bystander editing activities, causing limitations in their potential applications.

Off-target events of CBEs result from either the Cas9:sgRNA improperly targeting genomic loci, or the indiscriminate activity of the cytidine deaminase domain (3, 6–9). As such, the use of high-fidelity Cas9 variants and cytidine deaminase variants with enhanced DNA specificity have been reported to significantly decrease off-target events of CBEs (10–13). Additionally, a recent study showed that embedding the deaminase domain into the Cas9 gene, thus creating Cas-embedded CBE (CE_CBE), significantly minimized off-target editing in cell-based systems compared with linear N- or C-terminally linked fusions of Cas9 and deaminase (14). However, even these CE_CBE improvements cannot overcome some of the limitations due to the CBE delivery methods.

Genome editing tools, including CBEs, are primarily conveyed via plasmid delivery and have been known to increase the likelihood of off-target editing due to longevity of the editor within cells (10, 15, 16). Alternatively, direct electroporation of CBE:sgRNA ribonucleoprotein (RNP) complexes results in fast and efficient editing activity (10, 16–18) and the CBE is quickly degraded by endogenous cellular proteases (15, 19). Thus, CBEs delivered via electroporation reduce cellular off-target editing without compromising the on-target editing (20). In addition, RNP electroporation eliminates risks associated with undesired random DNA integration and unwanted immune responses due to excessive sgRNAs, thereby provide a safer alternative to plasmid delivery for base editing (19). However, the wider use of direct RNP delivery has been hampered due to the difficulty of producing high-quality, soluble, CBE or CE_CBE protein preparations needed (19). Thus, taken together, engineering highly soluble recombinant CE_CBE that can be used to produce CBE RNPs suitable for direct delivery would significantly improve safety, delivery, efficacy and activity of these complexes in a wide range of applications.

In this study, we take advantage of several existing technologies and combine them to engineer CE_CBE RNP systems that address the issues associated with off-target editing and direct delivery. Specifically, we designed the next-generation of highly soluble, stable CE_CBE (sCE_CBE) variants that can be expressed and purified in high yields. We showed that these variants exhibit highly efficient on-target editing, as well as shifted or expanded editing windows in human cells, thereby significantly improving editing efficacy and increasing target sequence space. In addition, we demonstrated that electroporating sCE_CBE RNPs together with uracil glycosylase inhibitor protein (UGI) in *trans* increased C-to-T editing efficiency and target purity in a dose-dependent manner, minimizing indel formation to untreated levels. These stabilized CE_CBE RNPs enabled us to use a single electroporation to effectively edit a therapeutically relevant locus for sickle cell disease in hematopoietic stem and progenitor cells (HSPCs). Significantly, the stabilized CE_CBE RNPs permitted for the transplantation of edited HSPCs confirming highly efficient editing in engrafted hematopoietic stem cells (HSCs) in mice. We have demonstrated that our engineered stabilized CE_CBEs overcome the limitations of off-target editing, RNP delivery, and restricted editing windows and are validated in a mammalian model. Thus our next-generation of CE_CBEs has a potential to safely and efficiently target disease-causing SNVs.

## Results

### Designing highly soluble and stable CE_CBEs

In designing our third-generation base editors (BE3s) for RNP delivery, we integrated several recent advances in the field. For the cytidine deaminase domain, we selected a single-domain cytosine base editor eA3A (N57G APOBEC3A). This variant has a strong TC motif preference, which minimizes sgRNA-independent and -dependent off-target DNA editing as well as off-target RNA editing activity (12). To generate a Cas9 embedded version of eA3A, we chose the 1048Thr-1063Ile region of nCas9 which was reported to tolerate domain insertion (14, 21, 22). The cytosine deaminase eA3A was embedded into the nCas9 by N- and C-terminal linkers. These linkers were optimized, as the linker length is known to affect the performance of base editors in terminally fused systems (23). We constructed and purified five CE_CBE with N and C linker lengths varying between 2 and 16 a.a. (CE_2_2_eA3A, CE_2_7_eA3A, CE_2_16_eA3A, CE_7_16_eA3A, and CE_16_16_eA3A) (**Fig. 1a**). Our AlphaFold modeling predicted that eA3A could be inserted into Cas9 and easily access the target ssDNA loop even with a short, 2 amino acid, Gly-Ser linker (**Fig 1b**). However, out of all the linkers tested (see Methods for details), eA3A flanked by 16 a.a. XTEN linkers on both sides (CE_16_eA3A) resulted in the highest protein and editing yields. In addition, nuclear localization signal (NLS) sequences that enhance editing efficiency (24–27) were added to CE_16_eA3A as three copies at the C-terminus and one copy at the N-terminus (**Fig. 1a**). Finally, the construct included UGI fusion at the C-terminus, as a way to decrease unwanted base substitutions (**Fig. 1a**). Thus, to develop the most stable and efficient CE_CBE, we optimally integrated these advances into our design.

**Figure 1.**
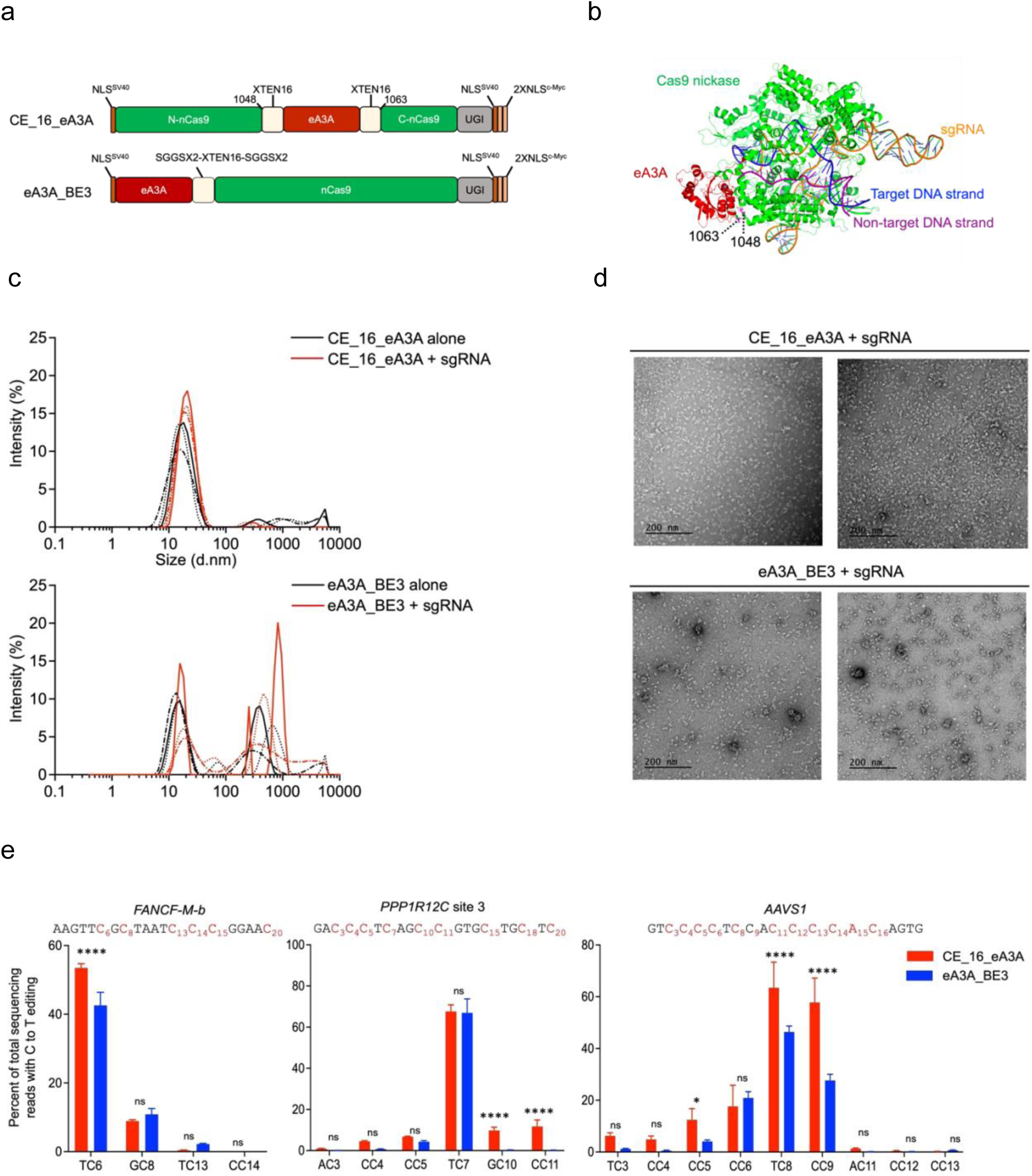
Engineered Cas-embedded (CE) eA3A proteins mediate efficient C to T conversion. (**a**) Schematic representation of eA3A-BE3 and the engineered base editors with 16 amino acid (a.a.) linker (CE_16_eA3A), where eA3A (red) was inserted into the 1048Thr-1063Ile region of nCas9 (Cas(D10A), green) via XTEN (16 a.a.) linker. For comparison, eA3A-BE3 was constructed by fusing eA3A to the N terminus of nCas9. (**b**) Model structure of CE_2_eA3A predicted by AlphaFold. (**c**) Dynamic light scattering (DLS) analysis (top) of CE_16_eA3A alone (black) and CE_16_eA3A RNPs(red), (prepared at a 1:1 sgRNA:protein ratio) – note the homogeneity and (bottom) of eA3A_BE3 alone(black) and eA3A_BE3 RNPs(red). Size distribution measured by DLS as a function of signal intensity of particles for n=3 independent measurements. (**d**) Representative TEM images (top) of CE_16_eA3A RNPs – note the mono-dispersion, (prepared at a 2:1 sgRNA:protein ratio) and (bottom) of eA3A_BE3 RNPs. Scale bars: 200 nm. (**e**) C to T editing frequencies of CE_16_eA3A and eA3A-BE3 after RNP delivery at *FANCF-M-b*, *PPP1R12C* site3, and *AAVS1.* Each target sequence is shown at the top. All editing frequencies were quantified by high-throughput sequencing (HTS) and are shown as means with error bars representing SD of n=3 biologically independent replicates on different days. Asterisks indicate statistically significant differences between eA3A-BE3 and CE_16_eA3A editing (****: *P* < 0.0001; *: *P* < 0.05; ns: *P >* 0.05, not significant). All statistical testing was performed using two-way ANOVA.

To benchmark our third-generation CE_CBE to a previously reported standard in the field (12), we also generated eA3A linked to the N-terminus of nCas9 with analogous NLS sequences, referred here as eA3A_BE3. We overexpressed CE_16_eA3A and eA3A_BE3 in *E. coli* and found that CE_16_eA3A showed enhanced expression, resulting in double the yield of purified protein ∼2 mg/L, compared to the N-terminally linked eA3A_BE3 (**Fig. S1**), and could be concentrated to 40 mg/mL. In fact, when we assessed our CE_16_eA3A by dynamic light scattering and electron microscopy, we observed that the protein forms a homogenous distribution of particles that shifts uniformly upon complexation with the sgRNA, in contrast to eA3A_BE3 which shows substantial levels of higher order aggregation (**Fig. 1c, d**). Due to the significantly improved stability, we named this system sCE_CBE for “stabilized Cas9-embedded cytosine base editor”. Taken together, our rationally engineered CE_16_eA3A variant represents the first highly soluble and homogeneous CE_CBE for use in direct delivery, and the first example of the sCE_CBE concept.

### High activity of stabilized Cas-embedded base editor RNPs

To confirm that we can achieve direct delivery of our highly purified sCE_CBEs without affecting activity, we targeted the *FANCF-M-b* site, *PPP1R12C* site 3, and *AAVS1* site with validated sgRNAs in HEK293T cells by electroporating the RNP complex (**Fig.1e**). We extracted genomic DNA and amplified the target genomic site for high-throughput sequencing, after three days. As expected, based on the known substrate specificity of A3A (28–30), for all the sites, both CE_16_eA3A RNPs and eA3A_BE3 RNPs preferentially edited cytosines in the TC motif context (TC> CC> GC> AC). High-throughput sequencing data showed that the C-to-T editing efficiencies of CE_16_eA3A RNPs were significantly higher for two of the three sites than that of eA3A_BE3 RNPs: *FANCF-M-b* TC_6_ (53.5±1.2% vs 42.6±3.8%, p < 0.0001), *AAVS1* TC_8_ (63.4±9.9% vs 46.4±2.3% p < 0.0001) and comparable (p= 0.7518) for the third *PPP1R12C* site 3 TC_7_ (**Fig. 1e**). Despite higher activity, the indel level formed by sCE_16_eA3A RNPs were similar or better, as in the case of *AAVS1,* than those of eA3A_BE3 (**Fig. S2**). We also tested the editing activity of other sCE_CBE constructs to examine whether varying the linker length between Cas9 and eA3A modulates efficiency of the CBE. All the variants tested displayed reduced efficiency compared to CE_16_eA3A, in agreement with our predictions and selection criteria. However, two of the constructs, CE_2_2_eA3A and CE_2_16_eA3A, while less, still had significant C to T editing activity (**Fig. S3**), and we included CE_2_2_eA3A (henceforth named CE_2_eA3A) in our further analysis. Taken together, these results demonstrate that CE_16_eA3A RNPs delivered via electroporation can effectively and accurately edit Cs in the TC motif context in human cells and suggest that linker length plays a role in modulating activity.

### Modulating the target of sCE_CBEs

All the base editors we constructed should preferentially target TC sites. To contrast editing activity and windows of CE_16_eA3A, CE_2_eA3A and eA3A_BE3 RNPs within different contexts, we analyzed average C-to-T base editing frequencies at 9 diverse target sites: *FANCF1* site 1, *VEGFA* site 2, *MSSK1-M-c*, *EMX1* site 1, *FANCF-M-b*, *RNF2*, *PPP1R12C* site 3, *PPP1R12C* site 6 and *AAVS1* (12, 14, 31) (**Fig. 2a, Table S1**). When combined, these target sites permit the interrogation of editing potential at target TCs from positions 2-3 through 19-20 (**Fig. 2b, Table S1a**) embedded in a variety of sequence contexts. These sequences also permitted the evaluation of bystander mutations CC, GC and AC (**Fig. S4, Table S1b**). We observed that CE_16_eA3A targets the TC most efficiently and specifically in each sequence at a total of ten sites, with eight being edited at more than 40% (**Fig. 2a**, **Table S1a**). Additionally, CE_16_eA3A exhibited only limited activity at six CC bystander sites, at one GC bystander site and no significant AC sites (**Fig. 2a, Table S1b**). This was further validated through amplicon sequencing of off-targeting sites of *VEGFA* site 2 (**Fig. S5**). CE_2_eA3A also edited shifted TC positions although less efficiently than CE_16_eA3A (**Table S1a**), while having better specificity (**Table S1b**). In contrast, eA3A_BE3 was overall both less efficient and less specific (**Table S1a, b**). Similarly, CE_16_eA3A RNPs transfected cells displayed negligible sgRNA-independent off-target C editing in an orthogonal R-loop assay at either site tested (**Fig. 2c, d**), despite better on-targeting (**Fig S6**); in contrast, eA3A_BE3 RNPs sgRNA-independent off-target C editing was above background. This contrast is much more dramatic when the plasmids encoding CE_16_eA3A and eA3A_BE3 were transfected. Notably, the RNP on-targeting control editing surpassed the plasmid editing (**Fig S6**). Overall, these results indicate that CE_2_eA3A and CE_16_eA3A have better specificity and higher efficiency than eA3A_BE3, and significantly expand the targets of the RNP-mediated CBE toolkit.

**Figure 2.**
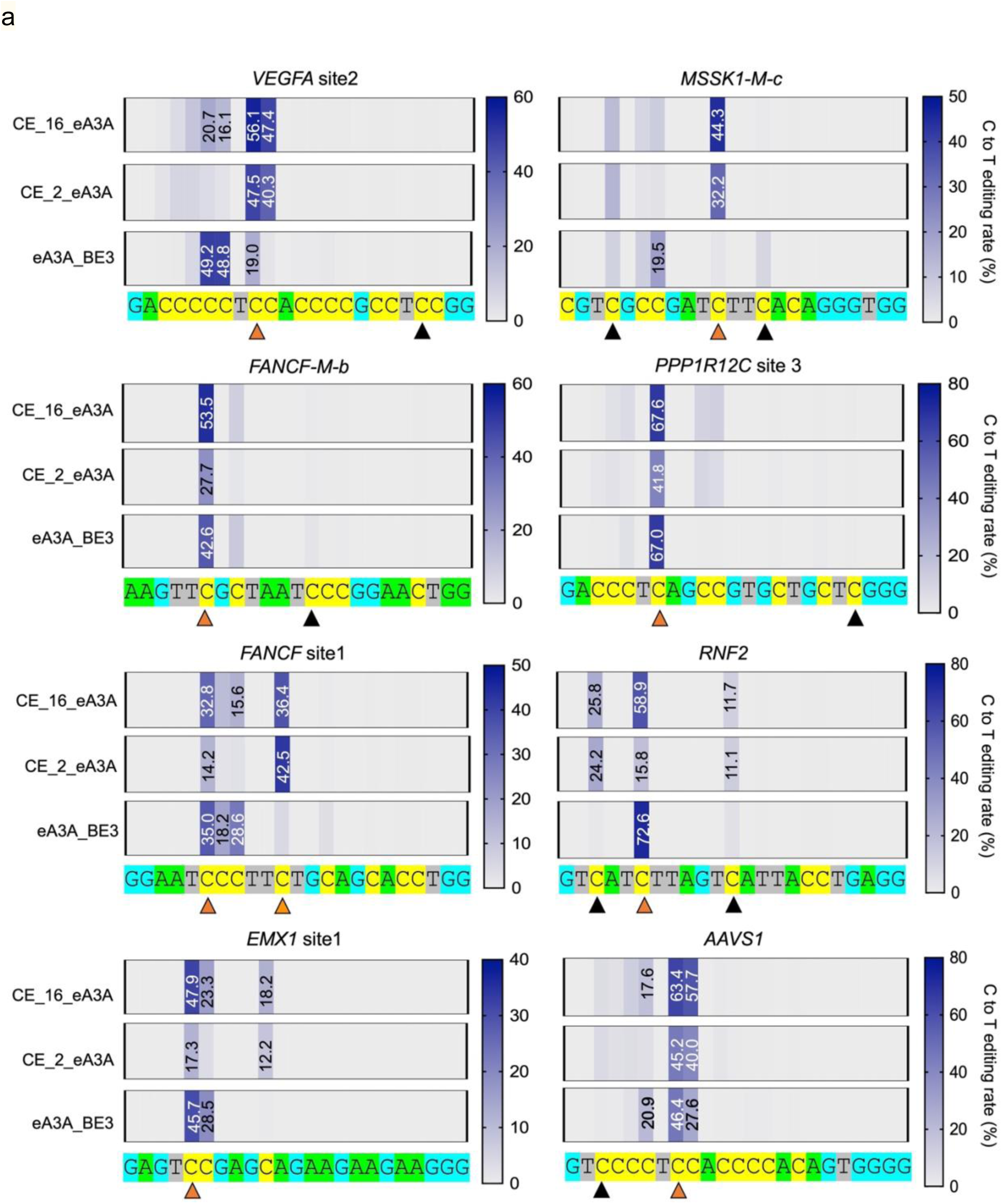
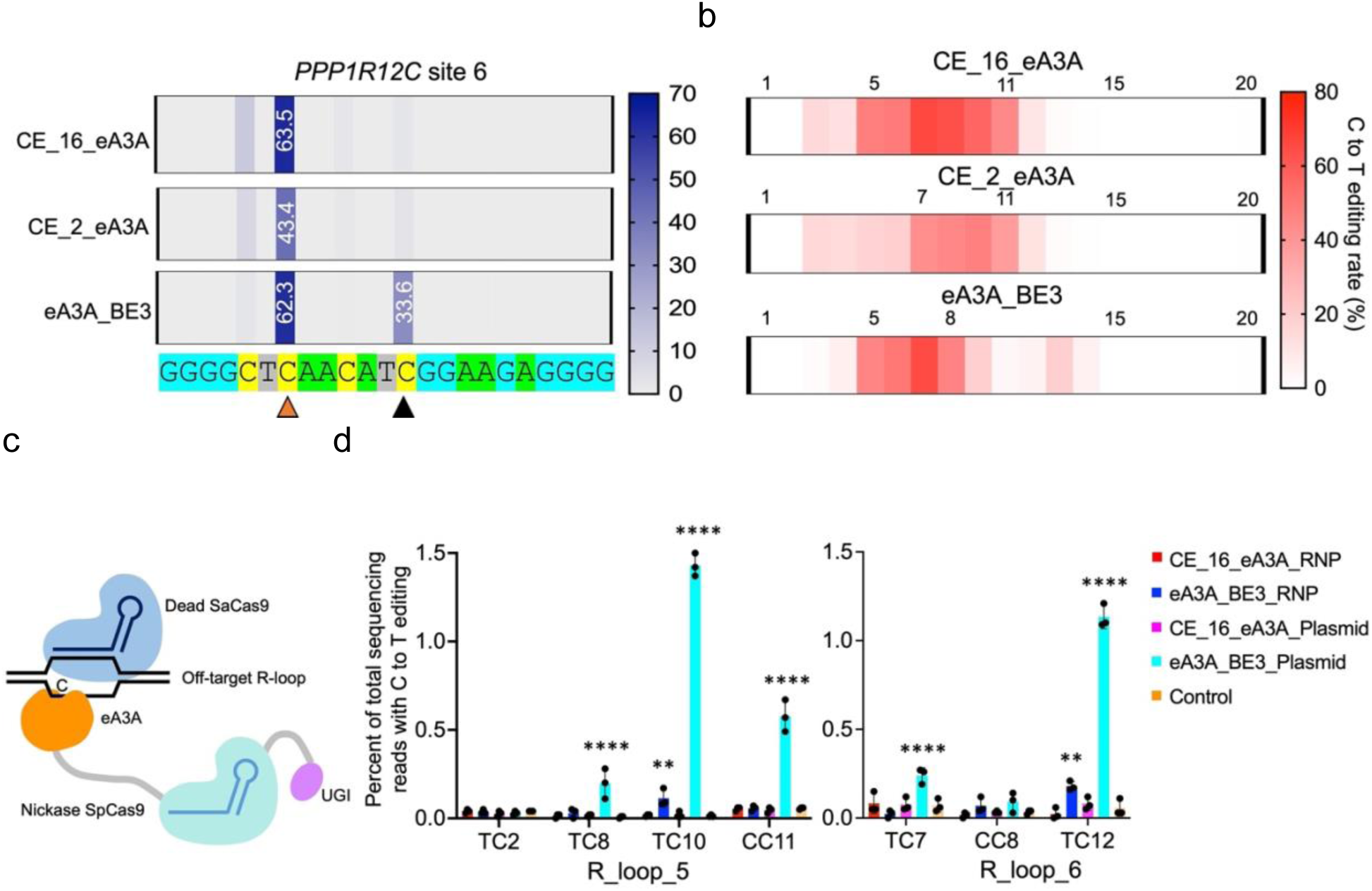
RNP-mediated on-target and off-target editing by CE_16_eA3A, CE_2_eA3A and eA3A-BE3. (**a**) Heat maps depicting the average C-to-T base editing frequencies of three independent replicates for CE_CBEs delivered as RNPs (4μM) at 9 target sites in HEK293T cells. Each target site in the editing window includes a cognate cytidine within a TC motif (marked with an orange arrow) and one or more bystander cytidines. Positions with a TC motif within the protospacers are indicated with triangles. (**b**) Schematic representation of the base editing window of CE_16_eA3A, CE_2_eA3A, and eA3A-BE3 with the heat map corresponding to the relative C to T editing activity at different positions of TC motifs within the protospacer regions. (**c**) Schematic representation of orthogonal R-loop assay. (**d**) sgRNA-independent off-target C to T editing frequencies of CE_16_eA3A (or eA3A_BE3) RNPs or plasmid detected by the orthogonal R-loop assay. Each R-loop was formed by cotransfection of base editor via RNPs or plasmid with SpCas9 sgRNA targeting *AAVS1* (on-target editing results presented in Fig. S6) and dSaCas9-encoding plasmid with SaCas9 sgRNA targeting R-loop 5 or 6. Data are shown as individual data points and mean±SD for n = 3 independent experiments. Asterisks indicate statistically significant differences in sgRNA-independent off-target C to T editing relative to untreated cells. (**: *P* < 0.01; ****: *P* < 0.0001). All statistical testing was performed using two-way ANOVA.

### Dose-dependent editing and target product purity of sCE_CBEs

In contrast to plasmid delivery, RNP delivery of gene editing systems provides a major advantage by allowing for a more limited temporal window of editing activity, as the protein-RNA complex gets quickly degraded. The advantage of this degradation is the reduction of cellular off-target editing compared to plasmid delivery where the editing system persists for a much longer time (20). We analyzed the dose-dependent editing frequencies at concentrations ranging from 1–10 μM of our sCE_CBEs after electroporation as *AAVS1* sgRNA complex (**Fig. 3a**). Since all three effectors edited TC7-8 at the *AAVS1* site efficiently in our previous assay, we compared the editing frequencies at this C8 site for each in HEK293T cells. The total base editing yield (C to any other base) of CE_16_eA3A reached its maximum (∼90%) at around 1 μM and plateaued from 1–10 μM whereas eA3A_BE3 saturated at 2 μM (**Fig. S7**). However, theC- to-T base editing efficiency, or product purity, increased dramatically from 28.6–38.9% to 55.5– 69.7% when the concentration of all tested RNPs was increased from 1 to 10 μM. At concentrations above 4 μM, CE_16_eA3A showed higher C-to-T base editing efficiency compared to eA3A_BE3 (*P*l<l0.05) (**Fig. 3a**). These results indicate that C-to-T base editing has a strong correlation with concentration of the base editors electroporated.

**Figure 3.**
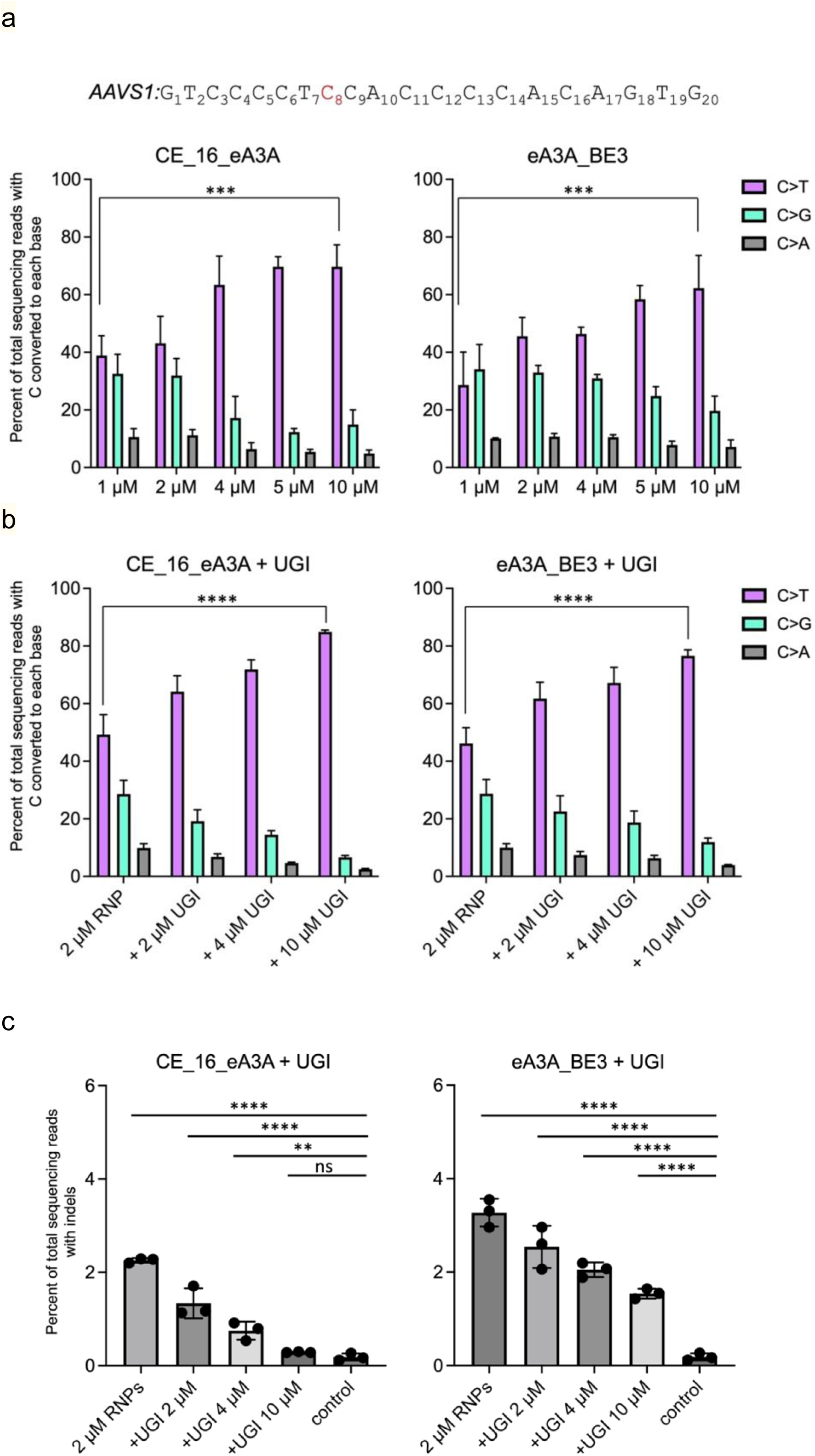
Dose-dependent editing yield, product purity, and indel formation. (**a**) Dose-dependent base editing of C8, which is preferentially edited by CE_16_eA3A or eA3A-BE3 at the *AAVS1* target site. Edited nucleotides that have been converted from target C8 are color coded; pink, C to T; light green, C to G; Gray, C to A. (**b**) C to T base editing purity and (**c**) indel formation after adding purified UGI proteins to RNPs of CE_16_eA3A or eA3A-BE3 at different ratios. Data are plotted as mean±SD, *n* = 3 independent experiments and analyzed using one-way ANOVA. Asterisks indicate statistically significant differences in editing efficiencies or indel levels observed between untreated cells and cells treated with RNPs of CE_16_eA3A or eA3A-BE3 and various ratio of purified UGI proteins. (ns: *P* > 0.05, not significant; **: *P* < 0.01; ***: *P* < 0.001; ****: *P* < 0.0001).

At 10 μM, all tested RNPs still produced some unwanted base substitutions (16.9±4.2 % C-to-G and 6.0±1.7 % C-to-A), demonstrating that the level of UGI was not sufficient to prevent these substitutions. To counter this, we investigated whether supplying recombinant UGI together with the sCE_CBE RNP *in trans* could decrease the unwanted substitutions. To this end, we purified UGI protein containing a nuclear localization signal (NLS) at the C-terminus (UGI-NLS), and electroporated HEK293T cells with 2 μM sCE_CBE RNPs and UGI-NLS protein in molar ratios of 1:1, 2:1, or 5:1 (UGI:CBE). Adding UGI-NLS protein to the RNPs significantly improved the editing purity in a dose-dependent manner (**Fig. 3b**). The addition of 5 molar equivalents of UGI-NLS protein to 2 μM CE_16_eA3A RNPs increased C to T editing from 43.1% to 85.0% (**Fig. 3b**), which is even higher than the C to T editing efficiency observed with 10 μM CE_16_eA3A RNP at the same site (70.0%). Similarly, the base editing product purity increased by adding UGI-NLS to RNPs for both eA3A_BE3 and CE_2_eA3A (**Fig. S8a,b**). Interestingly, adding UGI-NLS protein to RNPs also significantly decreased the formation of indels in a dose-dependent manner (**Fig. 3c, S8c**). Notably, indel formation was reduced to 0.28±0.15% for CE_16_eA3A and 0.35±0.32% for CE_2_eA3A, which is similar to background levels of untreated cells 0.19±0.08% when UGI-NLS protein was added to 2 μM CE_eA3A RNPs in 5:1 ratio. In contrast, eA3A_BE3 still caused considerable indel formation (1.54±0.11%) even in the presence of the UGI-NLS protein. These results suggest that the addition of UGI protein to stabilized Cas-embedded base editor RNPs at an optimal concentration can promote maximum C-to-T editing efficiency and purity while decreasing indel formation and unwanted base edits to background levels.

### Stabilized CE_CBE system achieves efficient base editing of a therapeutic target in human hematopoietic stem cells

To examine whether our new CE_16_eA3A RNP with improved solubility and activity could achieve high editing efficiency in human CD34+ HSPCs without inducing cytotoxicity, we targeted the +58 *BCL11A* erythroid enhancer region using a single electroporation (**Fig. 4**). Genetic disruption of this GATA1 motif within the enhancer sequence promotes therapeutic fetal hemoglobin (HbF) induction and can thereby ameliorate sickle cell disease and β-thalassemia (32, 33). We electroporated human CD34+ HSPCs with CE_16_eA3A or eA3A_BE3 RNPs containing the previously validated sgRNA-1620 (18) at concentrations varying from 5-50 μM (**Fig. 4**) (and similarly with CE_2_eA3A RNP with sgRNA-1618 (**Fig. S13**)). Previous studies required two cycles of electroporation to achieve high editing rates (∼90%) and resulted in significantly reduced HSPC viability (∼47%) and engraftment potential (18). In contrast, we detected little to no cellular toxicity at all tested concentrations of both CE_16_eA3A and eA3A_BE3 (**Fig. S9)**. As expected, both RNPs specifically edited the C6 position of the GATA binding motif with dose-dependent efficiency (**Fig. 4, S10)**. Notably, CE_16_eA3A RNPs achieved higher editing rates compared to eA3A_BE3 RNPs at all tested doses. At 5 μM, CE_16_eA3A exhibited 2.4-fold higher editing efficiency than eA3A_BE3 RNPs. At 20 μM, the CE_16_eA3A RNP produced 83.0±3.7% (C>T: 49.8±1.2%, C>G: 28.4±2.8%, C>A: 4.8±0.2%) base edits, which was better than eA3A_BE3 RNP (67.5.±8.7%) (**Fig. 4**), and A3A(N57Q)_BE3 RNP (63.6%) as previously reported (18). In fact, at 30 μM, CE_16_eA3A RNPs yielded 86.0±3.1% base edits without substantial cellular toxicity (viability 95.0±3.9%) through a single electroporation. The indel levels formed were not significant for either CE_16_eA3A RNPs or eA3A_BE3 RNPs at 5–50 μM (**Fig. S11)**, and neither RNPs induced any observable off-target editing at 58 of the 59 potential off-target sites in HSPCs (**Fig.S12**). At only one site (OT1), low-level C editing was observed in cells edited with CE_16_eA3A(3.7±1.1%) or eA3A_BE3 (1.3±0.4%) compared to negative control (0.4±0.2%) which are lower than the previously reported for A3A(N57Q)_BE3 (18). Thus, the sCE_CBE, CE_16_eA3A, successfully achieved very high base editing (86%), while limiting off-targeting and maintaining cellular viability.

**Figure 4.**
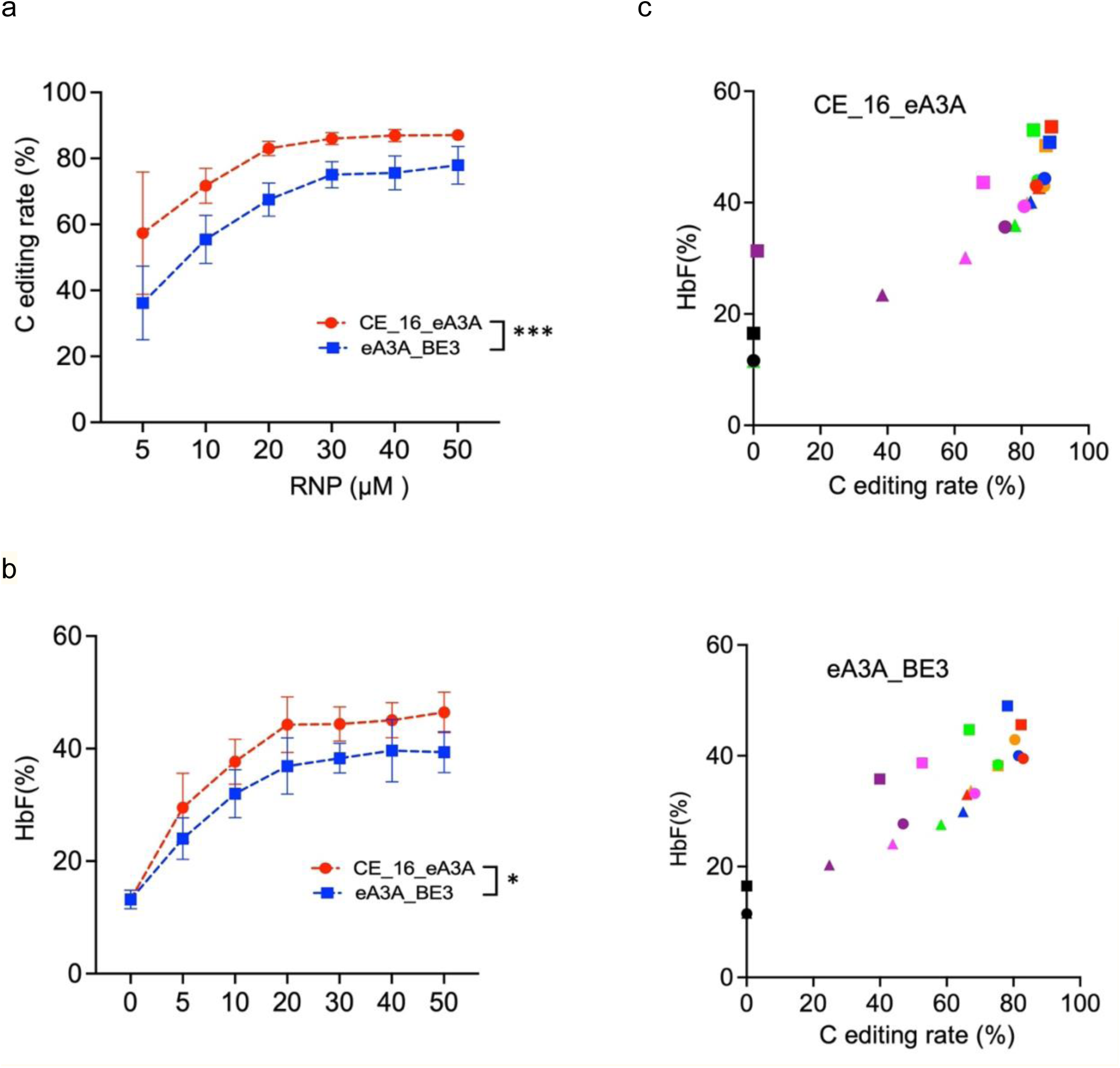
Dose dependent editing of *BCL11A* enhancer by CE_16_eA3A or eA3A_BE3 RNPs for HbF induction in CD34^+^ HSPCs. (**a**) Comparison of dose-dependent C base editing frequencies with CE_16_eA3A and eA3A_BE3 RNPs. Base edits quantified by high-throughput sequencing. (**b**) HbF levels of erythroid progeny after in vitro erythroid maturation from three healthy donors edited by different concentrations of CE_16_eA3A or eA3A_BE3 RNPs. HbF levels quantified by HPLC analysis. (**c**) Correlation of HbF levels versus C editing rates of erythroid progeny differentiated from CD34^+^ HSPCs edited by 5-50 μM CE_16_eA3A or eA3A_BE3 RNPs. Results were obtained from three independent experiments, presented as mean±SEM, and analyzed using two-way ANOVA. Asterisks indicate statistically significant differences in editing efficiencies and HbF levels observed between CE_16_eA3A RNPs and eA3A-BE3 RNPs. (*: *P* < 0.05; ***: *P* < 0.001). The various concentrations are shown with different colors. (Black: 0 μM, purple: 5 μM, Magenta: 10 μM, Green:20 μM, Orange: 30 μM, Blue: 40 μM, and Red: 50 μM)

Next, we used HbF production as a readout to demonstrate that RNP treatment with sCE_CBEs such as CE_16_eA3A can achieve a functional phenotype that correlated strongly with editing efficiency (**Fig. 4b, c**). At RNP concentrations above 30 μM, we were able to achieve 82.4–90.0% base editing by CE_16_eA3A that substantially elevated HbF protein levels by ∼3.5-fold, 35.9–54.6%, compared to the 13.2% (11.5–16.5%) baseline. Treatment with CE_16_eA3A resulted in slightly higher or similar HbF induction compared to eA3A_BE3 at all tested doses (**Fig 4b, c**). Taken together, these results confirmed that sCE_CBE RNPs delivered via electroporation are safe, effective and accurate tools for base editing in human cells.

### Stabilized CE_CBE direct delivery achieves efficient base editing in the context of stem cell xenotransplantation

To further investigate CE_16_eA3A editing in HSCs, we infused human CD34+ HSPCs unedited or edited with CE_16_eA3A-sg1620 RNPs from one healthy donor into NBSGW mice. After 16 weeks, the editing efficiency and human hematopoietic engraftment were evaluated from isolated bone marrow (BM). The overall base editing frequency in engrafted BM was 96.0±0.8%, which is significantly higher than the 70.3% previously reported after two cycles of A3A(N57Q) RNP electroporation (18) (**Fig. 5a**). We observed predominantly C to T (86.0±1.2%) base edits (5.5±1.9% to G and 4.5±1.7% to A). Flow cytometry analysis revealed ∼96% human cells in BM from both unedited (96.5±1.7%) and edited mice (96.9±2.7%) (**Fig. 5b**), as well as similar relative abundances of human B cells, myeloid cells, T cells, and HSPCs in mice transplanted. These results indicate that base editing with CE_16_eA3A RNPs did not affect the ability of transplanted HSCs to successfully engraft or differentiate into multiple lineages (**Fig. S14-15)**. In engrafting erythroid cells from CE_16_eA3A-edited HSPCs, the HbF level was markedly increased from 0% to 20.8% and the proportion of fetal cells was substantially increased from 5.0% to 59.8% (**Fig. 5c, d**). Collectively, our findings indicate that sCE_CBEs RNPs efficiently produce therapeutically relevant base edits in hematopoietic stem cells. Importantly, our findings suggest that sCE_CBEs offer significant improvements within a mammalian system over the existing CE_CBEs systems for which no *ex vivo* data has previously been reported.

**Figure 5.**
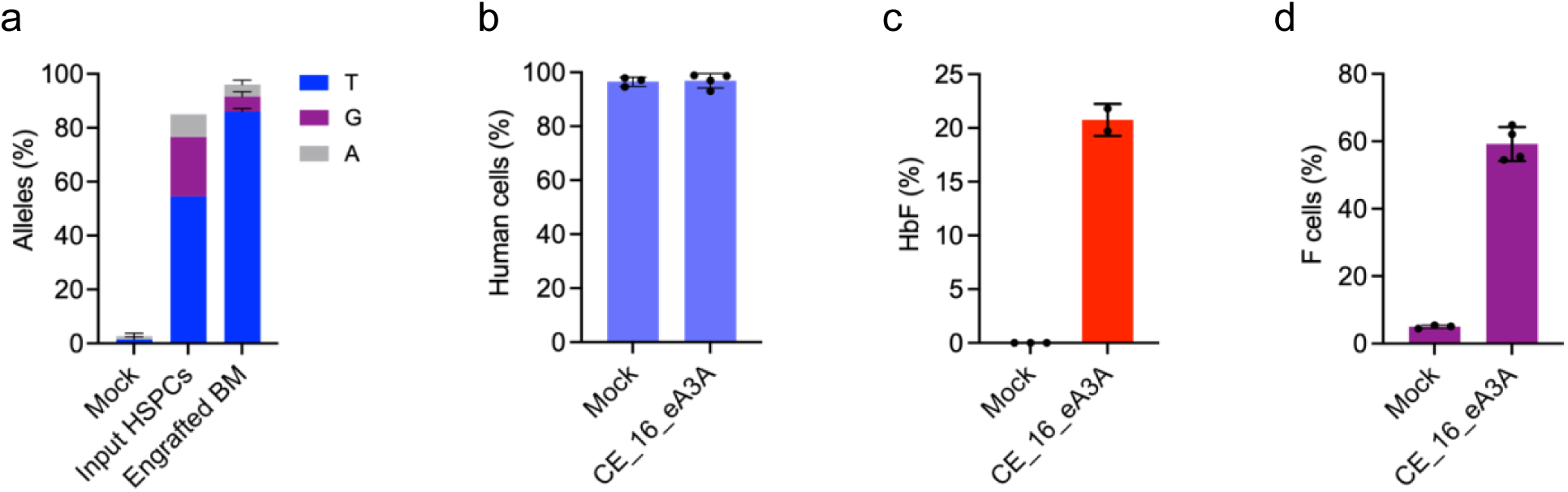
Highly efficient C to T editing in HSCs. Human CD34+ HSPCs from one healthy donor were electroporated with 30 μM of CE_16_eA3A RNPs. The treated HSPCs were infused into NBSGW mice and bone marrow (BM) was harvested 16 weeks after transplantation. (**a**) Base editing in unfractionated BM after 16 weeks as compared to input HSPCs. (**b**) Comparison of human chimerism in mice receiving unedited or edited HSPCs. Human cells in BM from all mice were quantified by flow cytometry using an anti-human CD45 antibody. Each dot indicates one mouse recipient. (**c**) HbF induction quantified by HPLC in human erythroid cells from engrafted BM. (**d**) Comparison of the percentage of fetal (F) cells between the mock and edited groups. The percentage of fetal cells in engrafted erythroid cells was measured by flow cytometry using an anti-human HbF antibody. Data are plotted as mean±SD (n=4 mice receiving edited cells and n=3 mice receiving unedited cells).

## Discussion

Currently, the main factors that limit the use of CBEs are the challenges associated with their delivery into the cells. Using plasmid delivery often results in persistent high concentrations of the base editor increasing the likelihood of off-target editing, while using direct delivery of RNPs often requires multiple rounds of electroporation to deliver sufficient amounts of base editor. In this context, repeated electroporation has been necessary because of poor biophysical properties of current CBEs. Therefore, despite offering more control over off-target effects, given the limited amount of base editor that it delivers and that repeated electroporation is damaging to cells, direct delivery of CBE RNPs has not been viable. Here, we addressed these challenges by designing stabilized Cas-embedded CBEs (sCE_CBEs) that can be produced in high yields and high purity. The resulting RNPs exhibited high editing efficiency, overcoming major limitations for the potential therapeutic utility and safety of these powerful editing tools. We also demonstrated that electroporating UGI-NLS in *trans* with these sCE_CBEs RNPs greatly improved C-to-T editing efficiency and product purity, and reduced the level of indel formation to background levels in a dose dependent manner.

Compared to previous attempts to directly deliver CBEs that required two cycles of electroporation and greatly compromised HSPC viability and engraftment potential (18), our CE_16_eA3A was able to achieve high editing efficiency through a single electroporation (up to 90% editing in HSPCs and 96% in bone morrow-repopulating HSCs). Importantly, directly delivered sCE_CBE RNP resulted in desired functional phenotype when used for genetic disruption of GATA1 motif associated with HbF production. Furthermore, we demonstrated that HSPCs edited via electroporated sCE_CBE RNP were fully compatible with xenotransplantation in NBSGW mice. Altogether, these results highlight safety, on-target activity and efficacy of our sCE_CBE system.

Our results have important implications for advancing base editors into the clinic. At the moment, two ongoing clinical trials (NCT03745287 and NCT03655678) are testing the use of autologous HSPCs modified with CRISPR-Cas9 for treatment of sickle cell disease and β-thalassemia (32, 33). Despite promising results, some questions remain regarding the safety of creating double-stranded breaks with the Cas9 effector (34). Based on our results, we suggest that the high editing efficiency and fidelity of our sCE_CBE, which does not generate double-stranded breaks or engage the DNA repair machinery to convert a single nucleotide, could provide an effective strategy for editing clinically relevant SNVs following a single RNP electroporation. Moreover, given the altered editing window of CE_eA3A RNPs, we expect that our tool will enable editing of Cs at otherwise inaccessible positions. Based on BEable-GPS (Base Editable prediction of Global Pathogenic-related SNVs) database (12, 35), we estimate that there are 229 SNVs correctable by CE_16_eA3A, as opposed to 143 correctable by eA3A_BE3. Therefore, sCE_CBEs could offer a strategy for expanding the target sequence space of base editors for future biomedical applications. We also envision that our approach will be applicable to building different context (C<u>C</u>, or G<u>C</u> or A<u>C</u>)-specific sCBE RNPs by switching eA3A with natural or modified/engineered domains of cytosine deaminases (31, 36), thereby further expanding the set of disease-relevant SNVs that can be targeted by high-fidelity base editors. Additionally, some of the more recently described base editors with more flexible editing windows, such as near-PAMless Cas9 enzymes (SpG and SpRY) and others (23, 31, 37, 38) may benefit from applying our approach to engineering variants for direct delivery to further improve editing efficacy and specificity. We anticipate that sCE_CBE editors, with improved solubility and enhanced on-target editing, will serve as promising agents for cytosine base editing at other disease-related sites in HSPCs and other cell types.

## Methods

### Plasmids and oligonucleotides

Sequences of CE_eA3A (or eA3A_BE3) proteins and expression plasmids used in this study are listed in the supplementary information 1. To construct pET21a-CE_2_eA3A, pET21a-CE_16_eA3A and pET21a-eA3A_BE3, we amplified eA3A, Cas9, and UGI genes from previously reported plasmids (addgene #131315) (12). We cloned the CBE constructs into the pET21a expression vector (Novagen) by standard molecular cloning methods. Supplementary 2 lists the sgRNA sequences and their corresponding target sites. For RNP delivery of CE proteins, chemically modified sgRNAs were designed to target sites and were synthesized by IDT Inc.

### Molecular Modeling

The structural model of CE_2_eA3A was predicted in AlphaFold 2 (39). We embedded the eA3A sequence at residue 1048-1063 of the SpCas9 protein sequence (PDB: 5F9R) (40) with GS (2 a.a.) linker. Additionally, we added a UGI sequence to the C-terminus of the SpCas9 protein. After prediction we added sgRNA and dsDNA to the model by merging PDB coordinates of our Cas-embedded eA3A protein with the sgRNA and dsDNA from a CRISPR-Cas9 complex (PDB: 5F9R). We optimized the structures using the Schrödinger Protein Preparation Wizard(41) at pH 8.0.

### Expression and purification of CE_2_eA3A, CE_16_eA3A and eA3A_BE3

For overexpression of CE_2_eA3A, CE_16_eA3A or eA3A_BE3 protein (42), the relevant plasmid was transformed into *E. coli* Rosetta2 (DE3) cells. The transformed cells were pre-cultured at 37 °C in LB medium with ampicillin (100 μg/mL) overnight, and the cell culture was inoculated into TB medium containing ampicillin (100 μg/mL), grown at 37 °C. When OD_600_ value reached ∼0.7-1.0, 0.5 mM IPTG was added to induce protein expression and the cell culture was further incubated overnight at 15°C. Cells were harvested by centrifugation at 5000 rpm for 20 min, resuspended in lysis buffer (50 mM Tris–HCl (pH 8.0), 1 M NaCl, 10 mM imidazole, 5% glycerol, 1 mM TCEP, EDTA-free protease inhibitor pellet (1 capsule/ 50ml, Roche)) and lysed by a cell disruptor. The lysate was clarified by ultracentrifugation at 18,000 rpm for 50 min. The supernatant was loaded onto a HisTrap FF column (GE Healthcare) equilibrated with lysis buffer, washed with washing buffer (50 mM Tris–HCl (pH 8.0), 1 M NaCl, 20 mM imidazole, 5% glycerol, 1 mM TCEP) and followed by elution using elution buffer (50 mM Tris–HCl (pH 7.5), 500 mM NaCl, 500 mM imidazole, 1 mM TCEP) with a linear gradient of imidazole ranging from 0 mM to 500 mM. The eluted proteins were dialyzed overnight at 4°C in 20 mM HEPES (pH 7.0), 150 mM NaCl, 10% glycerol, 1 mM TCEP. The dialyzed proteins were further purified using cation exchange column (5 mL HiTrap-S, Buffer A = 20 mM HEPES pH 7.0 +1 mM TCEP, Buffer B = 20 mM HEPES pH 7.0 + 1 M NaCl + 1 mM TCEP), and Superdex 200 size exclusion columns (running buffer A = 20 mM HEPES pH 7.5, 150 mM NaCl, 10% glycerol or running buffer B = 20 mM HEPES pH 7.5, 300 mM NaCl). The purified protein was concentrated using an Ultra-15 centrifugal filters ultracel-50K (Amicon) and flash frozen in liquid nitrogen. The size of purified proteins was measured by dynamic light scattering dispersed in PBS on a Zetasizer Nano ZS (Malvern Panalytical).

### Cell culture

Healthy human CD34 HSPCs were obtained from the Fred Hutchinson Cancer Research Center (Seattle, WA). Human CD34 HSPCs were thawed and cultured into serum-free medium Stem Cell Growth Medium (CellGenix, 20806-0500) supplemented with human Stem Cell Factor (SCF,100 ng/ml) (CellGenix, 1418-050), FMS-like Tyrosine Kinase 3 Ligand (Flt3L, 100 ng/ml) (CellGenix, 1415-050) and Thrombopoietin (TPO, 100 ng/ml) (CellGenix, 1417-050). After 48 hours of pre-stimulation, HSPCs were harvested for electroporation. Electroporated cells were cultured in erythroid differentiation medium (EDM) consisting of IMDM (GibcoTM, 12440061) supplemented with 330 µg/ml of Holo-Human Transferrin (Sigma-Aldrich, T0665-1G), 10 µg/ml of recombinant human insulin (Sigma-Aldrich, 19278-5ML), 2 IU/ml heparin (Sigma-Aldrich, H3149), 5% of human solvent detergent pooled plasma AB (Rhode Island Blood Center) and 3 IU/ml erythropoietin (AMGEM, 55513-144-10). During days 1-7 of culture, EDM was supplemented with 10-6M hydrocortisone (Sigma-Aldrich, H0135), 100 ng/ml human SCF (CellGenix, 1418-050) and 5 ng/ml of recombinant human IL-3 (PEPROTECH, 200-03). During days 7-11 of culture, EDM was supplemented with 100 ng/ml human SCF (CellGenix, 1418-050). During days 11-18 of culture, EDM had no additional supplements.

### RNP electroporation

To examine RNP-mediated genome editing in HEK293T cells, CE_2_eA3A, CE_16_eA3A or eA3A_BE3 protein was complexed with the corresponding sgRNA (supplementary information 2) in nuclease free water and incubated for 20 minutes at room temperature. Then, the resulting RNP complexes were mixed with HEK293T cells (1.0 × 10^5^) and electroporated using the neon transfection system.

For orthogonal R-loop assay with CE_16_eA3A (or eA3A_BE3) plasmid, 500 ng of dead SaCas9 plasmid (Addgene no. 138162), 200 ng of SaCas9 sgRNA plasmid, 500 ng of CE_16_eA3A (or eA3A_BE3) plasmid, 200 ng of AAVS1 sgRNA plasmid were cotransfected into HEK293T cells (4.0 × 10^4^) using 2 μl of Lipofectamine 3000 (catalog no. L3000008; Thermo Fisher Scientific). For orthogonal R-loop assay with base editor RNP, 500 ng of dead SaCas9 plasmid, 200 ng of SaCas9 sgRNA plasmid, and 700 ng of pUC19 plasmid (negative control plasmid) were cotransfected into HEK293T cells (4.0 × 10^4^) using 2 μl of Lipofectamine 3000. One day after transfection, cells treated without base editor plasmid were trypsinized and centrifuged for 5 min at 100xg. After resuspending the cells, the cells were electroporated with 75 pmole CE_16_eA3A protein (or eA3A_BE3) and 150 pmole sgRNA using the Neon Transfection System.

For editing the +58 *BCL11A* erythroid enhancer region in human CD34+ HSPCs, electroporation was performed using Lonza 4D Nucleofector. The RNP complex was prepared by mixing 100 pmol-1000 pmol of N-eA3A, NC16:CE-eA3A or NC2:CE-eA3A with 300 pmol-3000 pmol of sgRNA-1620 (IDT, TTTATCACAGGCTCCAGGAA) or sgRNA-1618 (IDT, TTGCTTTTATCACAGGCTCC), adding 2% of glycerol and P3 solution up to 10 µl. The RNP was incubated at room temperature for 10-15 min. 50,000-100,000 cells were suspended in 10 µl of P3 solution. The cell suspensions were mixed with RNP and transferred to cuvette (Lonza 4D, V4XP-3032) for electroporation with program EO-100. The P3 solution was removed after 15 min of incubation at room temperature. The electroporated cells were cultured in EDM.

### Base editing measurements

To determine on/off-target editing frequencies in HEK293T cells, cells were harvested 3 days after electroporation and genomic DNA was extracted with 100 μL lysis buffer (10 mM Tris-HCl, pH 7.5, 0.05% SDS, 25 μg/mL proteinase K (NEB)) at 37 °C with incubation for 1 hour. Proteinase K was inactivated by 30-minute incubation at 80 °C. The on- and off-target genomic sites (experimentally determined from a previous study) were PCR amplified with Phusion plus DNA polymerases (New England Biolabs) and locus-specific primers having tails complementary to the Truseq adapters: 98 degrees for 90 s; 30 cycles of 98 degrees for 15 s, 64 degrees for 30 s, and 72 degrees for 15 s; 72 degrees for 5 min. Resulting PCR products were subjected to Illumina deep sequencing. For deep sequencing, resulting PCR products were amplified with index-containing primers to reconstitute the TruSeq adapters using Phusion plus DNA polymerases (98l°C, 15ls; 62l°C, 30s; 72l°C, 20ls) ×10 cycles. Equal amounts of the PCR products from each experimental condition were pooled and gel purified. The purified library was deep sequenced using a paired-end 150lbp Illumina MiniSeq run. Frequencies of editing outcomes were quantified using CRISPResso2 (43). The percentage of editing was calculated as sequencing reads with the desired allele editing compared to all reads for the target locus.

To measure on-target editing frequencies of the +58 *BCL11A* erythroid enhancer region in human CD34+ HSPCs, cells were harvested 4-6 days after electroporation. Genomic DNA was isolated using the Blood and Tissue Kit (Qiagen, 69506) according to the vendor’s recommendations. The BCL11A enhancer DHS +58 on-target region was amplified with KOD Hot Start DNA Polymerase (EMD-Millipore, 71086-31) and corresponding primers (forward primer 5’-AGAGAGCCTTCCGAAAGAGG-3’ and reverse primer 5’ GCCAGAAAAGAGATATGGCATC-3’). The cycling conditions were 95 °C for 3 min; 30 cycles of 95 °C for 20 s, 60 °C for 10 s, and 70°C for 10 s; 70 °C for 5 min. A total of 1 µl of locus specific PCR product was used for indexing PCR using KOD Hot Start DNA Polymerase (EMD-Millipore, 71086-31) and TruSeq i5 and i7 indexing primers (Illumina) following the cycling conditions: 95 °C for 3 min; 10 cycles of 95 °C for 20 s, 60 °C for 10 s, and 70 °C for 10 s; 70 °C for 5 min. The indexed PCR products were evaluated by Qubit dsDNA HS Assay Kit (Thermo Fisher, Q32854), TapeStation with High Sensitivity D1000 Reagents (Agilent, 5067-5585) and High Sensitivity D1000 ScreenTape (Agilent, 5067-5584) and KAPA Universal qPCR Master Mix (KAPA Biosystems, KK4824/Roche 07960140001). The products were pooled as equimolar and subjected to deep sequencing using MiniSeq (Illumina).

To quantify off-target editing frequencies within human HSPCs edited by CE_16_eA3A or CE_2_eA3A or eA3A_BE3 RNPs, we performed amplicon deep sequencing of potential genomic off target sites in genomic DNA samples extracted from cells edited with CE_16_eA3A or CE_2_eA3A or eA3A_BE3 RNPs and from negative control cells. For sgRNA-1620, 59 potential genomic off target sites were identified previously (18). For sgRNA-1618, 146 potential off-target sites with three or fewer genomic mismatches and no bulges were identified using the CasOFFinder tool. Off-target sites were amplified with rhAmpSeq Library Mix 1 (IDT) and using rhAmpSeq forward and reverse assay primer pools. The cycling conditions were: 95 °C for 10 min; 14 cycles of 95°C for 15 s and 61°C for 8 min, and 99.5°C for 15 min. Locus specific PCR product was diluted to 1:20 and 11 µL was used for the indexing PCR with the cycling conditions: 95°C for 3 min; 24 cycles of 95°C for 15 s, 60°C for 30 s and 72°C for 30 s; and 72°C for 1 min. The resulting PCR products were evaluated by Qubit dsDNA HS Assay Kit (Thermo Fisher, Q32854), TapeStation with High Sensitivity D1000 Reagents (Agilent, 5067-5585) and High Sensitivity D1000 ScreenTape (Agilent, 5067-5584) and KAPA Universal qPCR Master Mix (KAPA Biosystems, KK4824/Roche 07960140001). The products were pooled as equimolar and subjected to deep sequencing using NovaSeq (Illumina).

### Hemoglobin HPLC

Erythroid cells were harvested 18 days after erythroid differentiation. Hemolysates were prepared by vertexing the cell pellet with Hemolysate reagent (Helena Laboratories, 5125). The hemolysates were mixed with D10 reagent (BioRad) and loaded on D10 Hemoglobin Testing System for the human hemoglobins.

### Human CD34+ HSPC transplant and engraftment analysis

NBSGW mice were infused with human CD34+ HSPCs by retro-orbital injection. Bone marrow (BM) was harvested 16 weeks after transplantation. For flow cytometry analysis, the BM was blocked with Human TruStain FcX™ (Biolegend, #422302) and TruStain fcX™ (anti-mouse CD16/32, Biolegend, # 101320) for 15 minutes at room temperature, followed by a 30-minute incubation on ice with Fixable Viability Dye (Thermo fisher, # 65-0865-18), mCD45 Monoclonal Antibody (Thermo fisher, # 61-0451-82), Mouse Anti-Human CD45 (BD Bioscience, # 560367), anti-human CD235a (BioLegend, # 349104), anti-human CD19 (BioLegend, # 302212), anti-human CD33 (BioLegend, # 366608), anti-human CD34 (BioLegend, # 343504), and anti-human CD3 (BioLegend, # 300420). Samples were acquired and recorded on a BD FACSAria II and data were analyzed with FlowJo™ software.

For MACS hCD235a isolation, BM was incubated with hCD235a microbeads (Miltenyi, # 130-050-501) for 15 minutes on ice, followed by LS column hCD235a positive selection (Miltenyi, # 130-042-401). 20% of the hCD235a+ cells were analyzed for HbF content by flow cytometry. The cells were stained with Hoechst 33342 for 20 minutes at 37 °C, fixed with 0.05% glutaraldehyde (Sigma, #G6257) in PBS at room temperature for 10 minutes, then permeabilized with 0.1% Triton X-100 (Life, # HFH10). After permeabilization, cells were stained with mouse-antihuman CD235a (BioLegend, #349115) and HbF (Life, #MHFH014) in 0.1% BSA/PBS for 30 minutes on ice. Samples were acquired and recorded on a BD FACSAria II and data were analyzed with FlowJo™ software. 80% of the hCD235+ cells were also analyzed for hemoglobin content by HPLC. The cells were lysed with hemolysate reagent (Helena Laboratories cat# 5125) and HPLC analysis was performed with D-10 Hemoglobin Analyzer (Bio-Rad).

## Acknowledgements

This work was supported by National Institutes of Health R01AI150478 (previously R01GM118474). We also acknowledge Dr. Ala Shaqra for laboratory support and Drs. Milka Kostic and April April Pawluk for editorial support and suggestions from LifeScienceEditors.com.

