## Supplemental Figures and Tables for "Direct delivery of stabilized Cas-embedded base editors achieves efficient and accurate editing of clinically relevant targets"

*Joint last author

[1]Department of Biochemistry and Molecular Biotechnology, UMass Chan Medical School, Worcester, MA 01605, USA

[2]Department of Molecular, Cell and Cancer Biology; University of Massachusetts Chan Medical School, Worcester, MA 01605, USA

[3]Division of Hematology/Oncology, Boston Children's Hospital, Boston, MA, USA

[4]Department of Pediatric Oncology, Dana-Farber Cancer Institute, Boston, MA, USA

[5]Department of Pediatrics, Harvard Stem Cell Institute, Broad Institute of Harvard and MIT, Harvard Medical School, Boston, MA, USA


| | Page |
|---|---|
| Supporting Figures | S2-S19 |
| Supporting tables | S20-S21 |
| Protein sequences | S22-S23 |
| gRNA sequences | S24 |
| Expression plasmids used in this study | S24 |

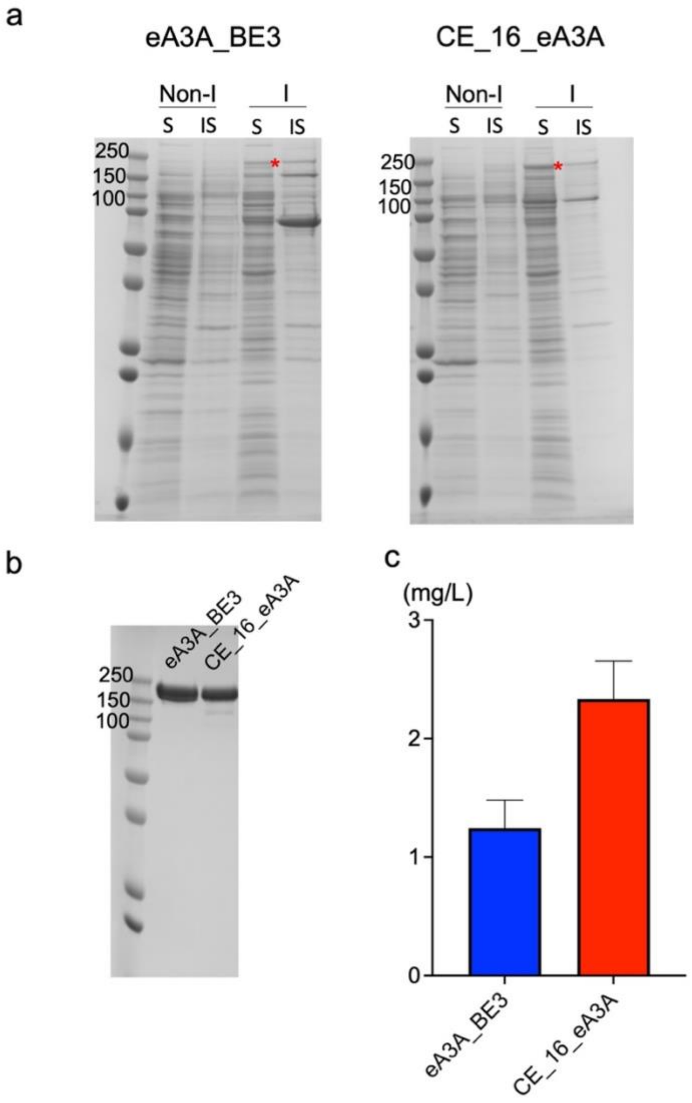


**Figure S1**. Purification of CE_16_eA3A and eA3A-BE3. (a) Coomassie-stained SDS-PAGE showing expression levels of CE_16_eA3A or eA3A-BE3 protein in *E. coli*, in the soluble (S) and insoluble (IS) fractions. (b) Coomassie-stained SDS-PAGE showing eluates of CE_16_eA3A or eA3A-BE3 protein purified by nickel affinity, mono S and Q ion exchange, and size exclusion columns next to protein lysates. The purity of purified proteins was determined using SDS-PAGE and gel staining. The purity of CE_16_eA3A or eA3A-BE3 protein was estimated to be ~ 97-99 %. (c) The yield of purified CE_16_eA3A or eA3A-BE3 protein. Bars represent mean values, and error bars represent the SD of three independent replicates.

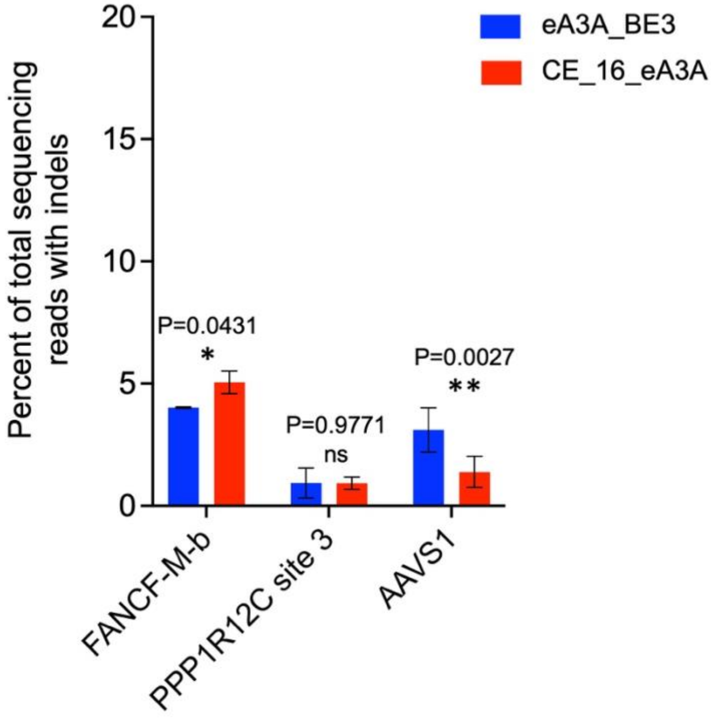


**Figure S2**. Indel formation by CE_16_eA3A or eA3A-BE3 RNPs. Indel levels of CE_16_eA3A or eA3A-BE3 after RNP delivery at 3 target sites (*FANCF-M-b*, *PPP1R12C* site6, and *AAVS1*). Indel levels represent the sum of all indel values in the editing window at each target site. Asterisks indicate statistically significant differences in indel levels observed between eA3A-BE3 and CE_16_e3A at each site (ns: $P > 0.05$, not significant; *: $P < 0.05$; **: $P < 0.01$). All statistical testing was performed using two-way ANOVA.

a

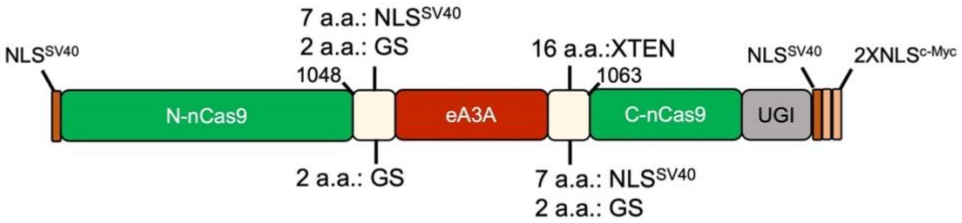


b

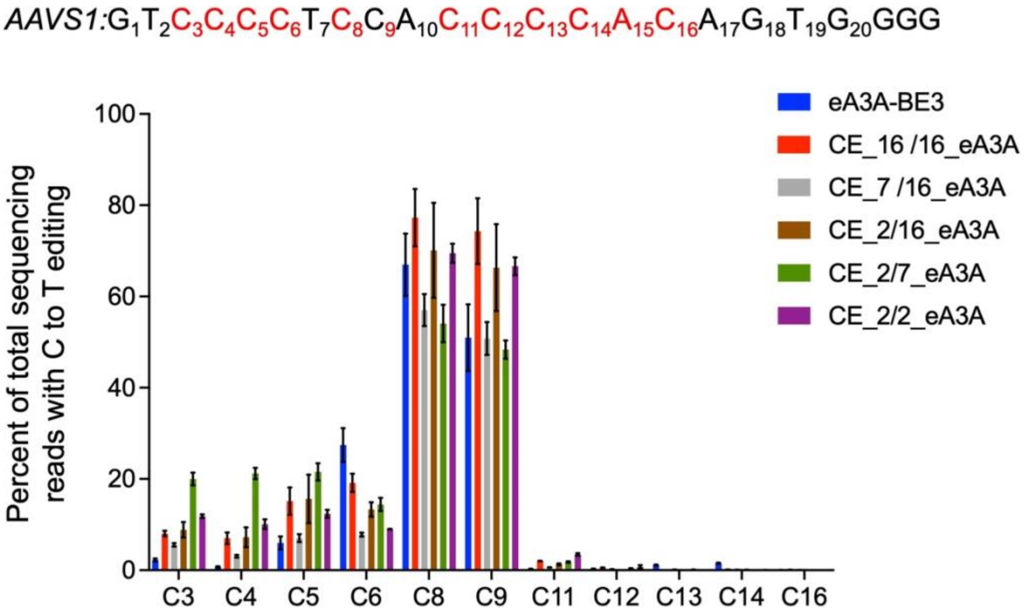


**Figure S3**. Base editing activities of engineered CE-eA3A variants with various linkers. (a) The schematic representation of engineered CE-eA3A variants with varying linker lengths between 16 a.a. and 2 a.a. (b) The frequencies of C to T editing by CE-eA3A variants after RNP delivery at *AAVS1*.

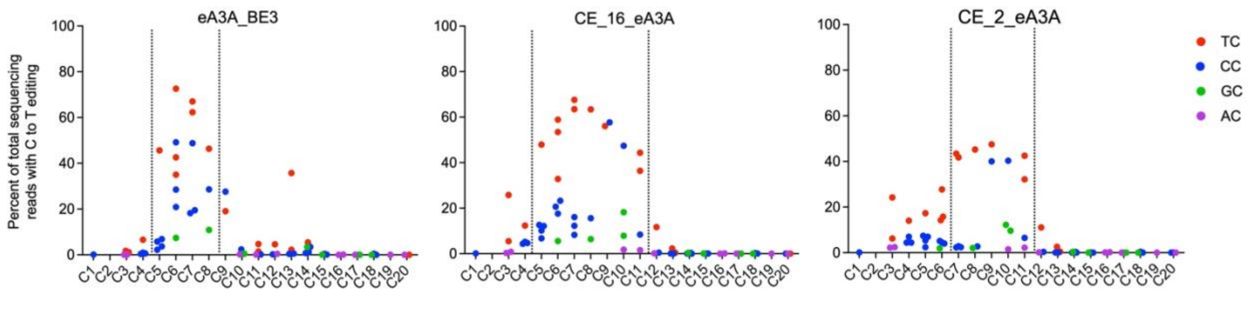


**Figure S4**. Average C-to-T base editing efficiencies of CE_16_eA3A, CE_2_eA3A, and eA3A-BE3 in different contexts (T<u>C</u> (red), C<u>C</u> (blue), G<u>C</u> (green) and A<u>C</u> (magenta)) from 9 target sites. Each data point represents the mean of triplicate measurements for each C in each target site.

a b

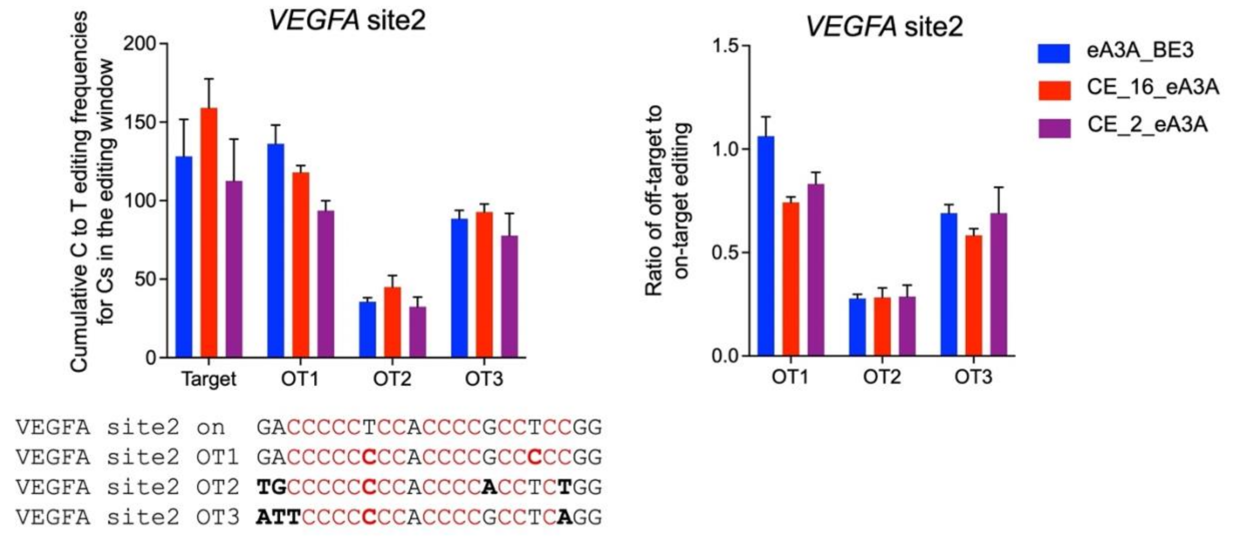


**Figure S5**. Editing at off-target sites of *VEGFA site2* by CE_16_eA3A, CE_2_eA3A, or eA3A-BE3 RNPs. (a) Each on/off- targeting efficiency was calculated by accumulating of all edited Cs in the editing window of on-target site or off-target sites. (b) The ratio of off-target to on-target C to T editing efficiencies. Bars show mean values and error bars the SD from three independent replicates.

a

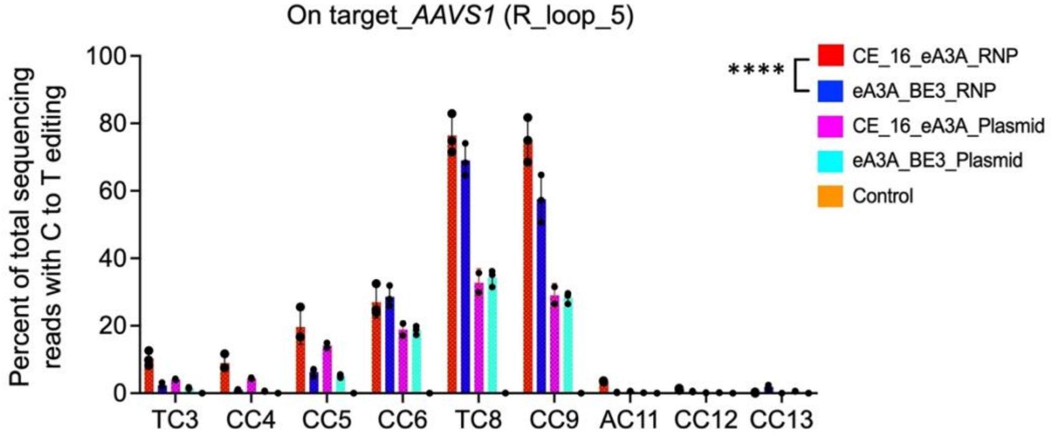


b

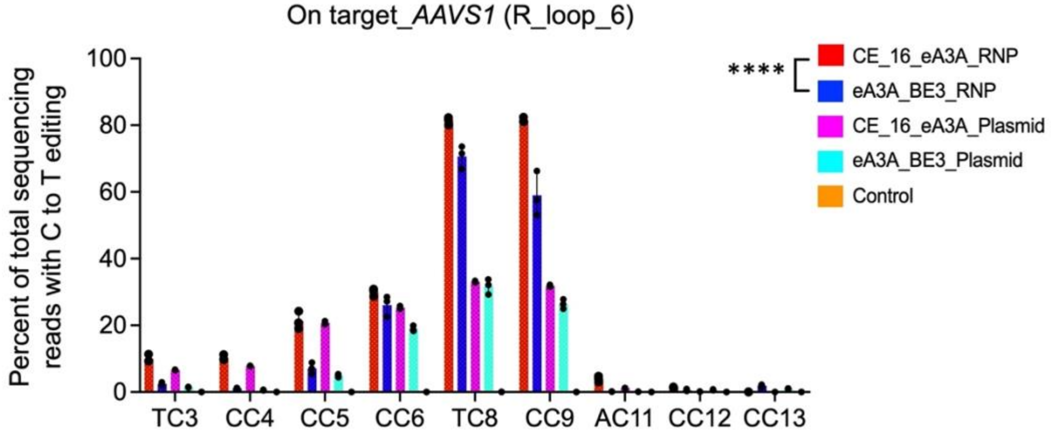


**Figure S6.** On-target control editing frequencies at *AAVS1* locus to test sgRNA-independent off-target editing by CE_16_eA3A (or eA3A_BE3) RNPs (7.5μM) or plasmid at SaCas9 R loop_5 (a) and 6 (b). Data are shown as individual data points and mean±SD for n = 3 independent experiments. Asterisks indicate statistically significant differences in the cumulative C to T editing efficiency observed between CE_16_eA3A RNPs and eA3A-BE3 RNPs (****: $P < 0.0001$). All statistical testing was performed using two-way ANOVA.

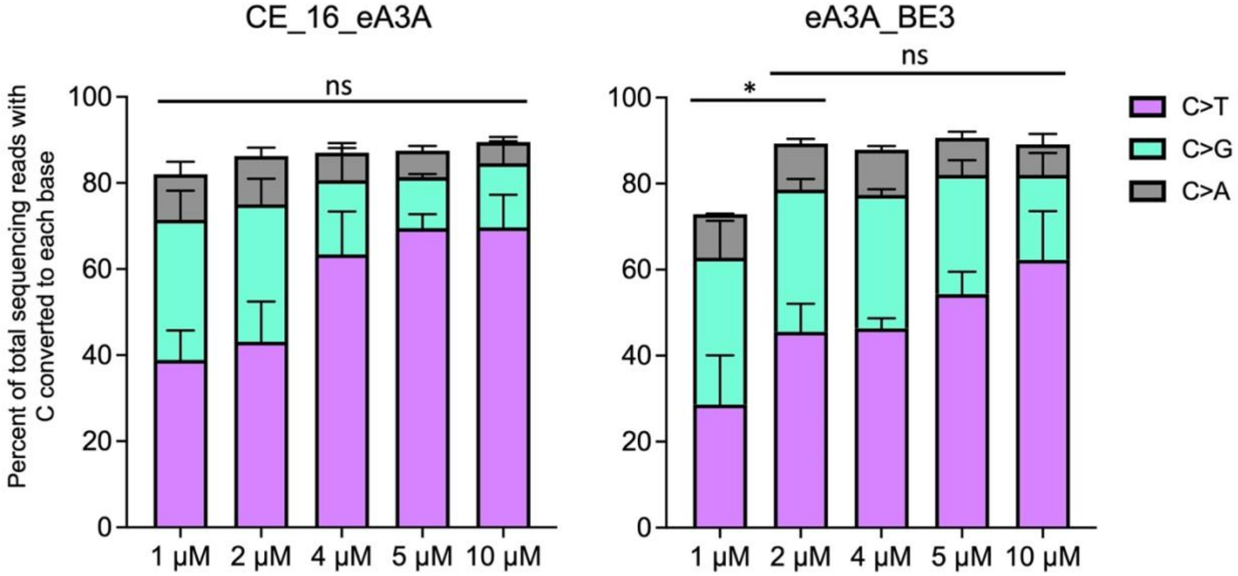


**Figure S7**. The total base editing yield (C to any other base) of CE_16_eA3A and eA3A-BE3.

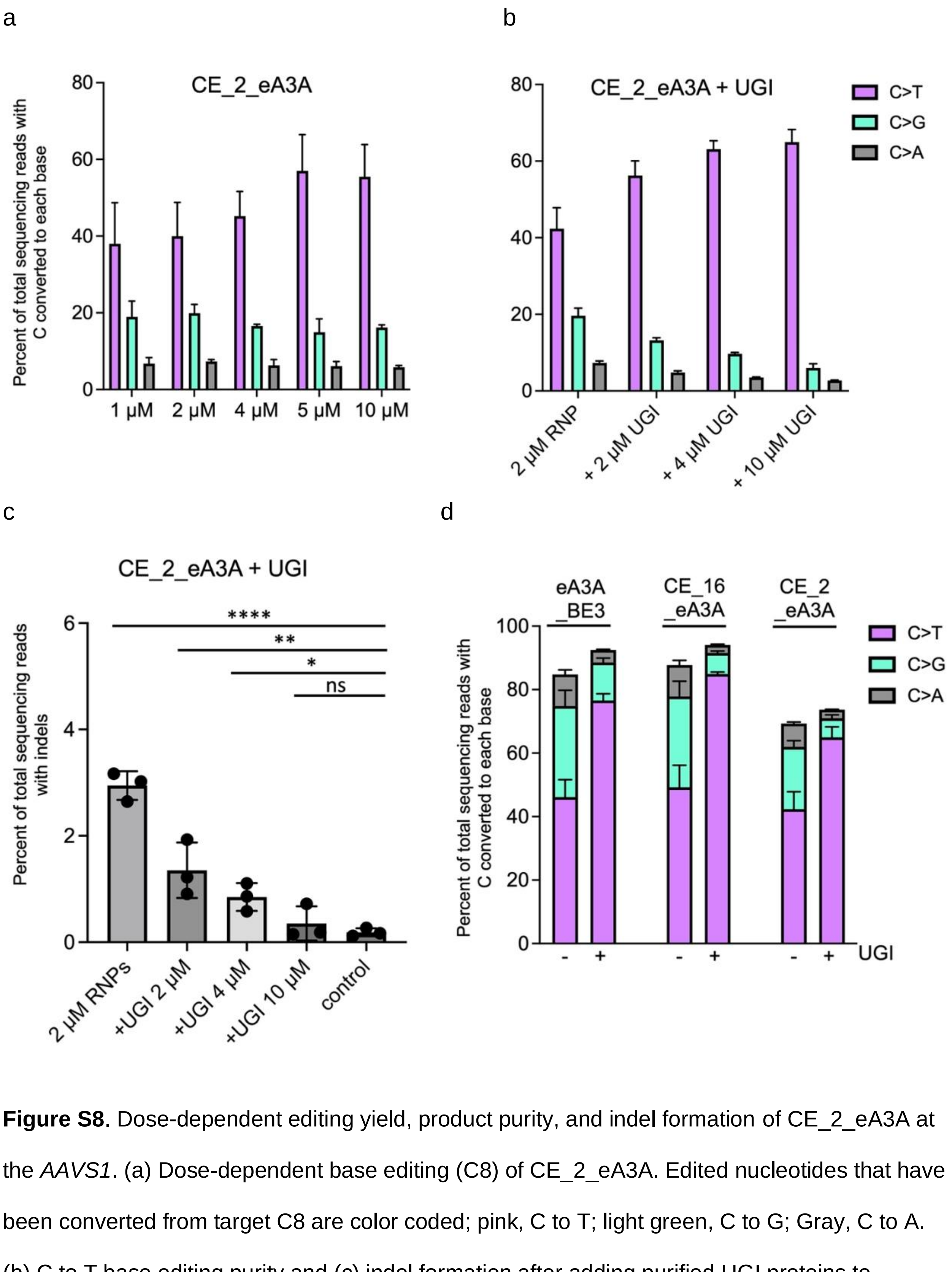


**Figure S8**. Dose-dependent editing yield, product purity, and indel formation of CE_2_eA3A at the *AAVS1*. (a) Dose-dependent base editing (C8) of CE_2_eA3A. Edited nucleotides that have been converted from target C8 are color coded; pink, C to T; light green, C to G; Gray, C to A. (b) C to T base editing purity and (c) indel formation after adding purified UGI proteins to

CE_2_eA3A RNPs at different ratios. (d) Comparison of product purity with/without adding purified UGI proteins to CE_16_eA3A, CE_2_eA3A, or eA3A-BE3 RNPs. Asterisks indicate statistically significant differences in indel levels observed between untreated cells and cells treated with RNPs of CE_16_eA3A, or eA3A-BE3 and various ratio of purified UGI proteins. (ns: $P > 0.05$, not significant; *: $P < 0.05$; **: $P < 0.01$; ****: $P < 0.0001$). All statistical testing was performed using one-way ANOVA.

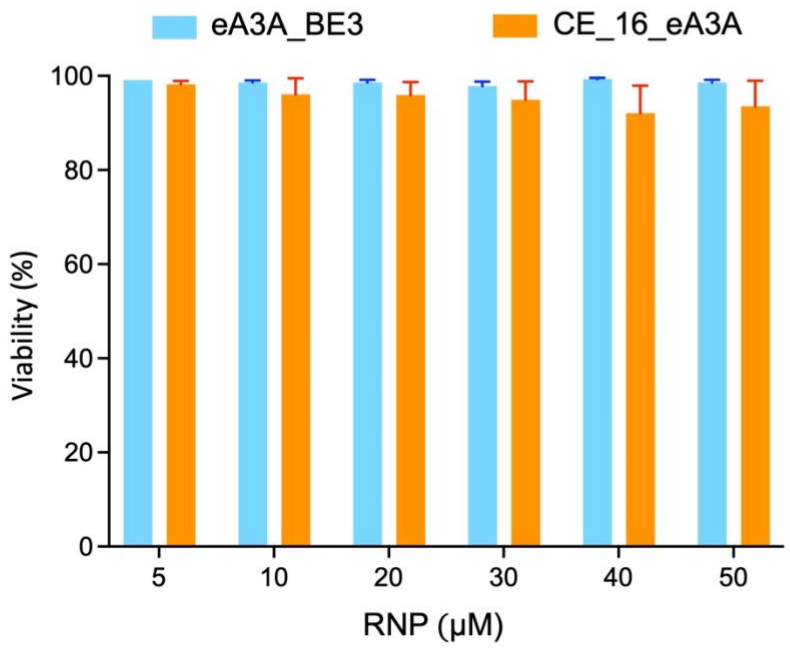


**Figure S9**. Comparison of viability of CD34$^+$ HSPCs edited with various concentrations (5–50 μM) of CE_16_eA3A or eA3A-BE3 RNPs.

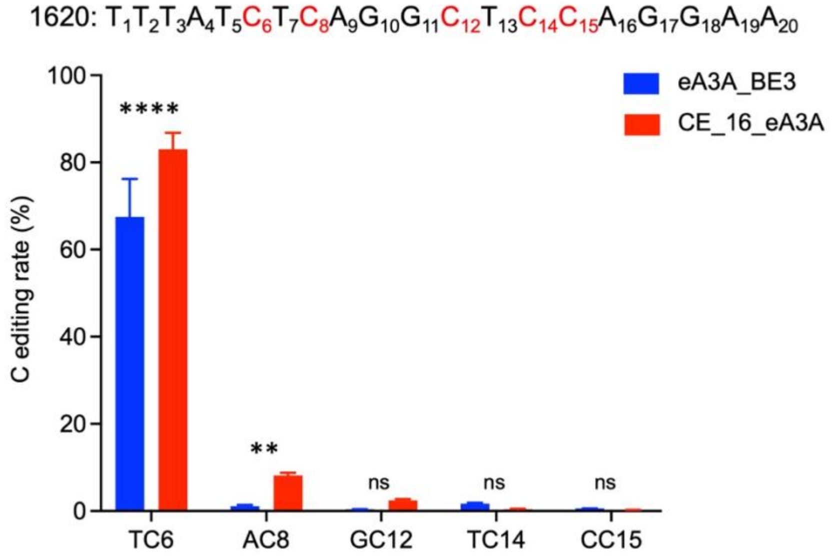


**Figure S10**. Comparison of C editing rate of CE_16_eA3A and eA3A_BE3 (20 μM) complexed with 1620 gRNA in human CD34+ HSPCs from three independent healthy donors. C editing rates were measured with deep sequence analysis. Asterisks indicate statistically significant differences between CE_16_eA3A and eA3A-BE3 editing (****: $P < 0.0001$; **: $P < 0.01$; ns: $P > 0.05$, not significant). All statistical testing was performed using two-way ANOVA.

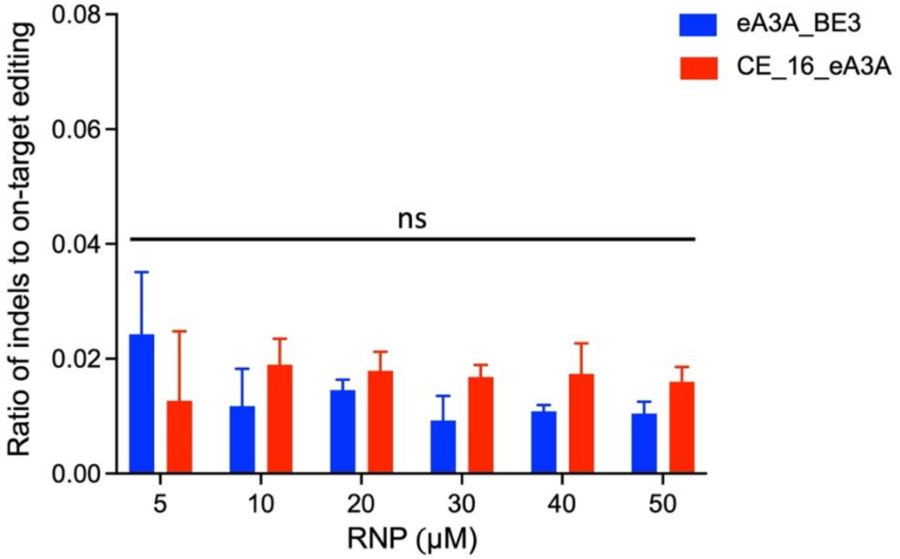


**Figure S11**. Indel formation by different concentrations (5-50 μM) of CE_16_eA3A or eA3A-BE3 RNPs at the +58 *BCL11A* erythroid enhancer region in human CD34+ HSPCs. There were no statistically significant differences in indel levels observed between eA3A-BE3 and CE_16_e3A at each concentration (ns: $P > 0.05$, not significant). All statistical testing was performed using two-way ANOVA.

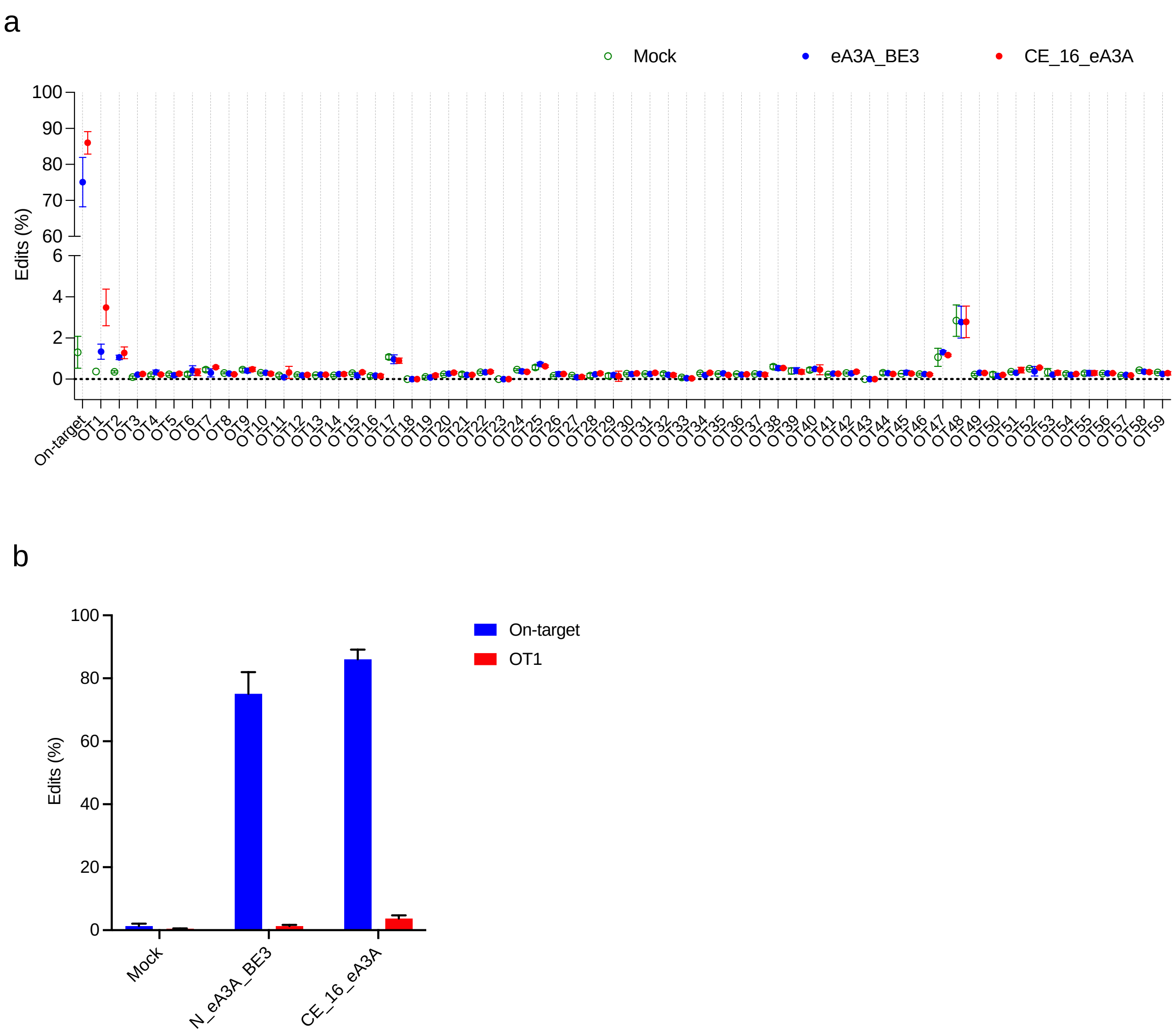


**Figure S12.** Guide RNA(1620)-dependent off-target editing analysis of HSPCs with CE_16_eA3A and eA3A-bE3 RNPs. (a) High-throughput sequencing data of 59 potential off-target sites within CD34$^+$ HSPCs edited with 30 μM CE_16_eA3A RNPs (red) and eA3A-BE3 RNPs (blue). Negative controls (mock) are indicated by green circles. (b) Comparison of on-target (blue) and off-target (red) C editing efficiency at OT1 by CE_16_eA3A RNPs and eA3A-bE3 RNPs.

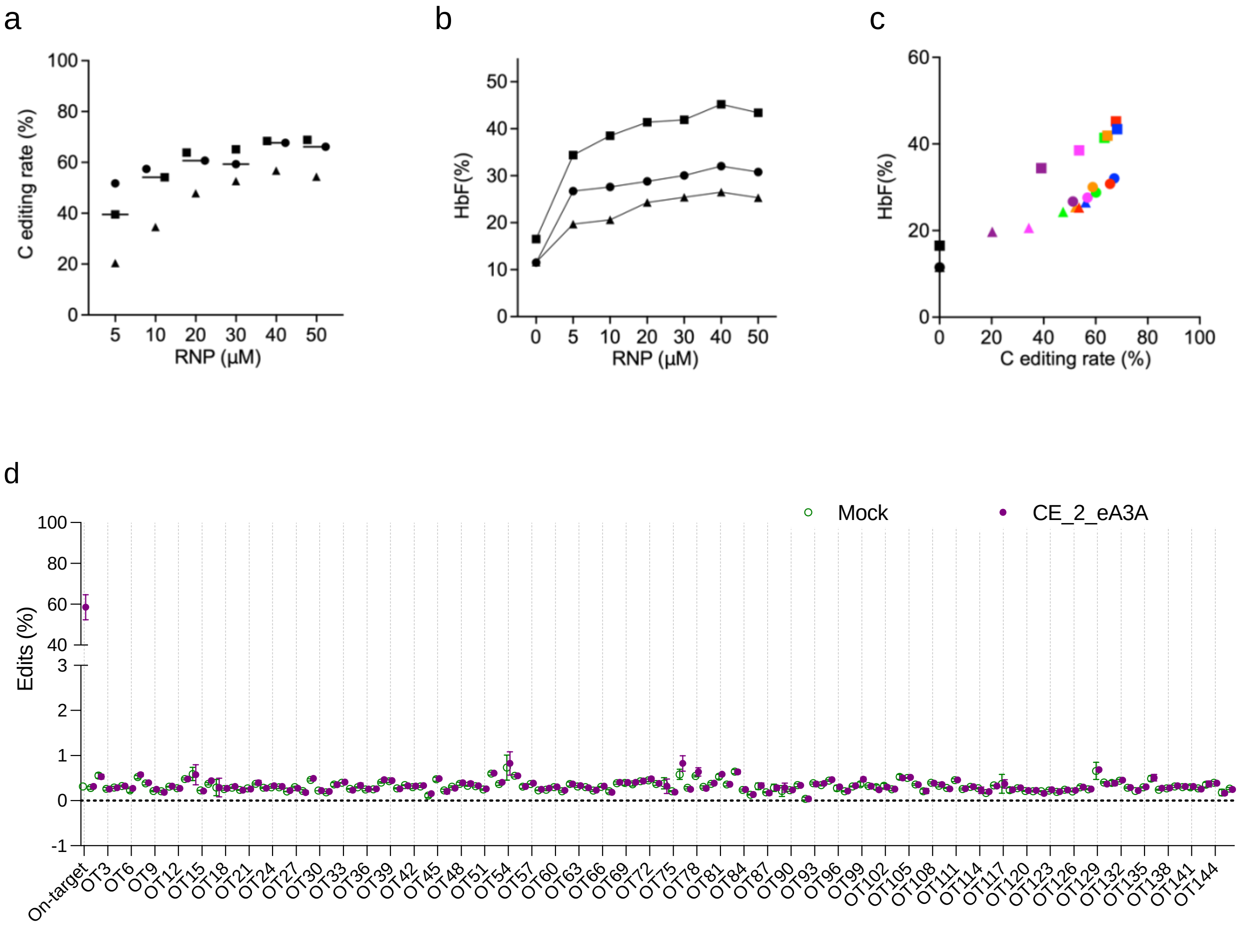


**Figure S13.** On/off-target editing analysis of HSPCs edited with CE_2_eA3A RNPs. The *BCL11A* enhancer region was targeted with CE_2_eA3A RNPs with sgRNA-1618 to edit position C11 of the GATA binding motif, to test whether the shifted editing window can be leveraged to eliminate off-target editing. (a) Dose dependent editing rates with CE_2_eA3A coupled with sgRNA-1618 targeting *BCL11A* enhancer in $CD34^+$ HSPCs by high-throughput sequencing analysis. (b) HbF levels by HPLC analysis of erythroid progeny after dose-response of CE_2_eA3A RNPs electroporation of HSPCs. (c) Correlation of HbF levels versus C editing rates in erythroid cells differentiated from CE_2_eA3A RNP-edited $CD34^+$ HSPCs. Circles, squares, or triangles are indicated in an individual healthy donor. Mean values are indicated in bars. The various concentrations are shown with different colors. (Black: 0 μM, purple: 5 μM, Magenta: 10 μM, Green:20 μM, Orange: 30 μM, Blue: 40 μM, and Red: 50 μM) (d) Using the

CasOFFinder tool, 146 potential genomic off target sites with 3 or fewer mismatches relative to the on-target *BCL11A* enhancer sequence were identified. In human $CD34^+$ HSPCs edited by 30 μM CE_2_eA3A RNPs (purple), the C editing efficiency of each of these 146 sites was quantified by high-throughput sequencing. Negative controls (mock) are indicated by green circles. CE_2_eA3A RNPs effectively produced ~59.0 % base edits, where A3A(N57Q)-BE3 RNP showed almost no activity under the same conditions with this sgRNA (Zeng, et al. *Nature Medicine* 26, 535-541, 2020). At all examined off sites, there was no detectable editing by CE_2_eA3A RNPs relative to the negative control, indicating that CE_2_eA3A RNPs effectively eliminated off-target editing while achieving efficient editing at the desired location.

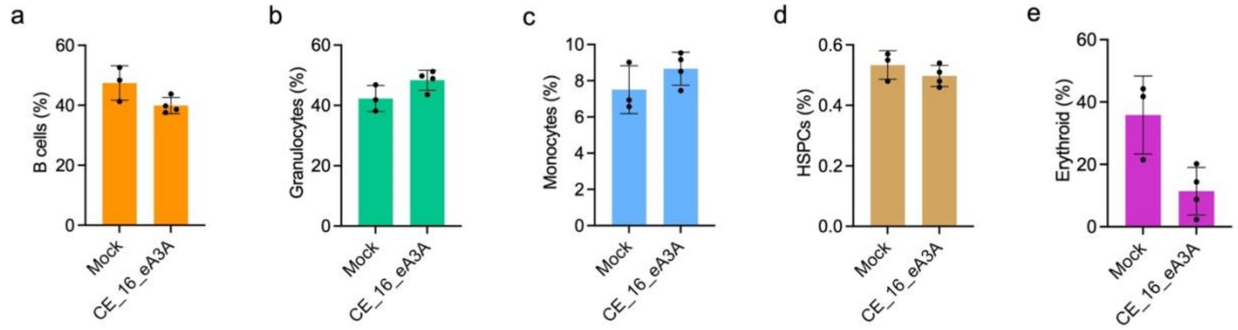


**Figure S14**. Percentage of B cells, myeloid, erythroid, and HSPC human lineages 16 weeks after transplantation with unedited or edited HSPCs. Data are plotted as mean±SD (n=4 mice receiving edited cells and n=3 mice receiving unedited cells).

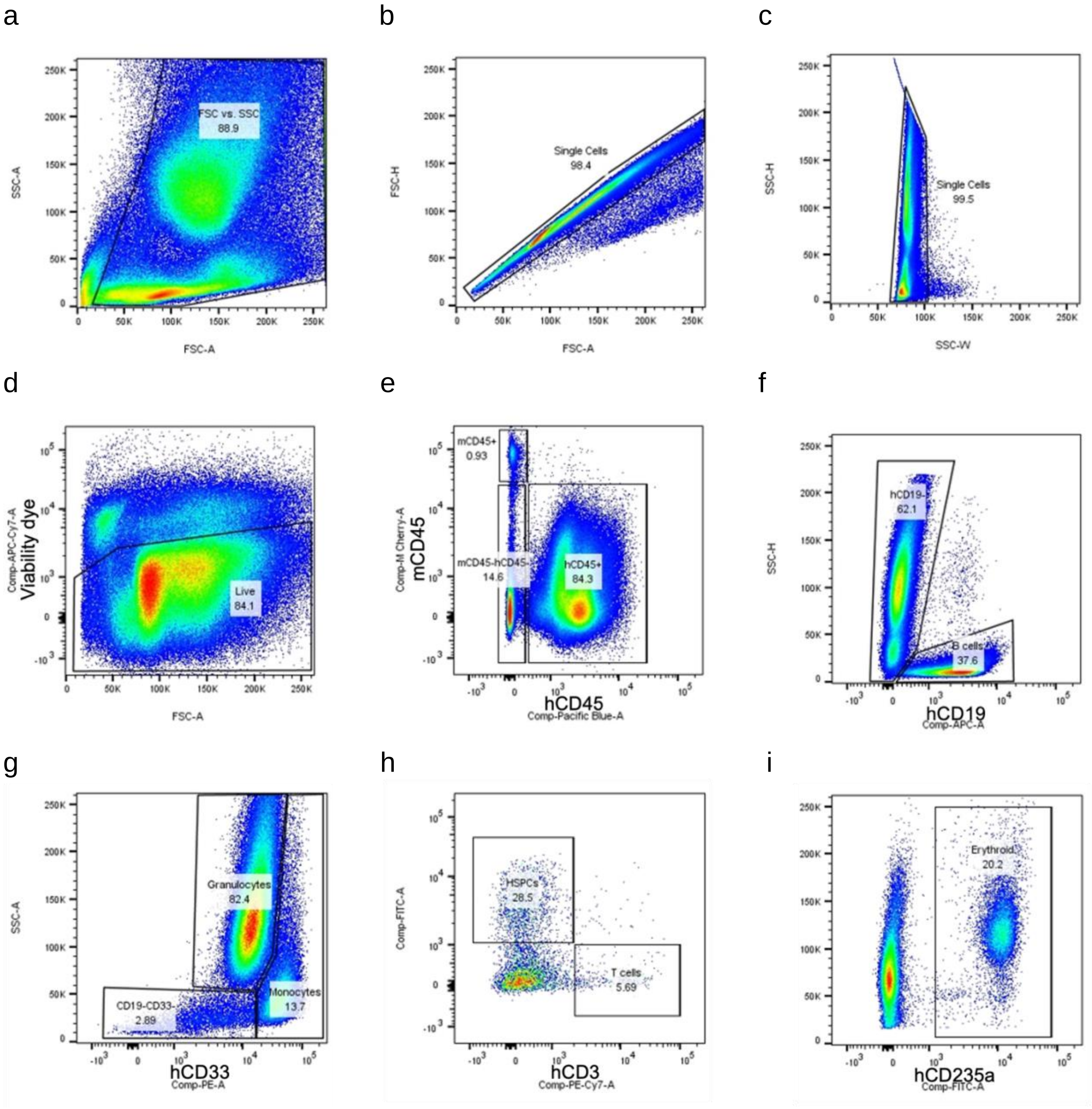


**Figure S15.** Representative flow cytometry analysis of xenografted bone marrow. (a–d) Live cells from engrafted mouse BM. (e) Human cells gated from hCD45+ population, mouse cells gated from mCD45+ population. (f) B cells gated from hCD45+CD19+ population. (g) Granulocytes gated from hCD45+CD19-CD33dim with SSC high population. Monocytes gated from hCD45+CD19-CD33+ with SSC low population. (h) CD34+Lin-(HSPCs) gated from hCD45+CD19-CD33-CD34+ population. T cells gated from hCD45+CD19-CD33-CD3+ population. (i) Erythroid cells gated from hCD45-mCD45- hCD235a+ population.

**Table S1a.** Average C-to-T base editing efficiencies of CE_16_eA3A, CE_2_eA3A, and eA3A-BE3 at a cognate cytidine in a TC motif within the editing window.

| | Average C to T base editing efficiencies | | |
|---|---|---|---|
| Cognate C within the editing window | CE_16_eA3A | CE_2_eA3A | eA3A_BE3 |
| *VEGFA* site2 TC9 | 56.1±6.2 | 47.5±9.6 | 19.0±4.5 |
| *MSSK1-M-c* TC11 | 44.3±3.8 | 32.2±7.0 | 1.5±0.1 |
| *FANCF-M-b* TC6 | 53.5±1.2 | 27.7±1.8 | 42.6±3.8 |
| *PPP1R12C* site3 TC7 | 67.6±3.3 | 41.8±3.6 | 67.0±6.8 |
| *FANCF* site1 TC6 | 32.8±2.0 | 14.2±2.7 | 35.0±3.4 |
| *FANCF* site1 TC11 | 36.4±1.4 | 42.5±1.8 | 4.7±0.3 |
| *RNF2* TC6 | 58.9±8.6 | 15.8±3.9 | 72.6±8.1 |
| *EMX1* site1 TC5 | 47.9±4.6 | 17.3±2.8 | 45.7±6.5 |
| *AAVS1* TC8 | 63.4±9.9 | 45.2±6.4 | 46.4±2.3 |
| *PPP1R12C* site6 TC7 | 63.5±3.2 | 43.4±6.6 | 62.3±3.2 |

**Table S1b.** Average C-to-T base editing efficiencies of CE_16_eA3A, CE_2_eA3A, and eA3A-BE3 for bystander cytidines (within the editing window but not within a TC motif), and for cytidines outside the editing window. (ND: not detectable)

| | Average C to T base editing efficiencies | | |
|---|---|---|---|
| Bystander C or C outside editing window | CE_16_eA3A | CE_2_eA3A | eA3A_BE3 |
| *VEGFA* site2 AC3 | 0.8±0.1 | 2.4±0.3 | ND |
| *VEGFA* site2 CC4 | 5.2±0.6 | 7.0±1.5 | 0.6±1.5 |
| *VEGFA* site2 CC5 | 12.2±1.1 | 7.0±1.7 | 5.8±1.4 |
| *VEGFA* site2 CC6 | 20.7±2.4 | 5.1±1.4 | 49.2±8.1 |
| *VEGFA* site2 CC7 | 16.1±2.2 | 2.8±1.4 | 48.8±8.8 |
| *VEGFA* site2 CC10 | 47.4±6.5 | 40.3±10.6 | 2.3±0.7 |
| *MSSK1-M-c* TC4 | 12.4±0.7 | 14.0±4.0 | 6.6±0.3 |
| *MSSK1-M-c* GC6 | 5.5±0.6 | 2.0±0.5 | 7.4±0.5 |
| *MSSK1-M-c* CC7 | 8.2±0.5 | 2.4±0.2 | 19.5±1.0 |
| *MSSK1-M-c* TC14 | 0.3±0 | 0.2±0 | 5.5±0.3 |
| *FANCF-M-b* GC8 | 8.9±0.4 | 2.1±0.2 | 10.9±1.7 |
| *FANCF-M-b* TC13 | 0.4±0.1 | 0.4±0.1 | 2.2±0.2 |
| *PPP1R12C* site3 CC4 | 4.5±0.3 | 4.0±0.4 | 0.7±0.1 |
| *PPP1R12C* site3 CC5 | 6.7±0.3 | 2.0±0 | 4.3±0.5 |
| *PPP1R12C* site3 GC10 | 9.8±1.6 | 8.8±0.7 | 0.3±0 |
| *PPP1R12C* site3 CC11 | 11.7±3.2 | 6.8±0.8 | 0.2±0.1 |
| *FANCF site1* CC7 | 12.3±1.3 | 2.3±0.4 | 18.2±3.0 |
| *FANCF site1* CC8 | 15.6±1.5 | 2.7±0.3 | 28.6±2.9 |
| *FANCF site1* GC14 | 0.4±0 | 0.4±0.1 | 3.5±0.2 |
| *RNF2* TC3 | 25.8±3.5 | 24.2±3.2 | 1.6±0.3 |
| *RNF2* TC12 | 11.7±3.6 | 11.1±2.3 | 4.6±0.6 |
| *EMX1* site1 CC6 | 23.3±3.3 | 4.5±0.9 | 28.5±5.6 |
| *EMX1* site1 GC10 | 18.2±1.6 | 12.2±2.4 | 0.5±0.1 |
| *AAVS1* TC3 | 6.2±1.2 | 7.0±0.6 | 1.2±0.2 |
| *AAVS1* CC4 | 4.9±1.3 | 4.9±0.8 | 0.5±0.2 |
| *AAVS1* CC5 | 12.4±4.4 | 5.8±1.2 | 4.1±0.6 |
| *AAVS1* CC6 | 17.6±8.2 | 5.1±2.0 | 20.9±2.4 |
| *AAVS1* CC9 | 57.7±9.5 | 40.0±6.4 | 27.6±2.4 |
| *AAVS1* AC11 | 1.3±0.3 | 2.1±0.1 | 0.1±0 |
| *PPP1R12C* site6 GC5 | 12.7±0.4 | 7.4±1.1 | 2.2±0.2 |
| *PPP1R12C* site6 AC10 | 1.8±0.3 | 1.5±0.3 | 0.2±0.1 |
| *PPP1R12C* site6 TC13 | 2.3±0.3 | 2.6±0.7 | 33.6±1.9 |

**Protein amino acid and sgRNA sequences, and expression plasmids**

1) Protein amino acid sequences

eA3A-BE3

MPKKKRKVTSPGPREASPASGPRHLMDPHIFTSNFNNGIGRHKTYLCYEVERLDNGTSVKMDQHRGFLHG
QAKNLLCGFYGRHAELRFLDLVPSLQLDPAQIYRVTWFISWSPCFSWGCAGEVRAFLQENTHVRLRIFAA
RIYDYDPLYKEALQMLRDAGAQVSIMTYDEFKHCWDTFVDHQGCPFQPWDGLDEHSQALSGRLRAILQNQ
GNIDEFSGGSSGGSSGSETPGTSESATPESSGGSSGGSPWDKKYSIGLAIGTNSVGWAVITDEYKVPSKK
FKVLGNTDRHSIKKNLIGALLFDSGETAEATRLKRTARRRYTRRKNRICYLQEIFSNEMAKVDDSFFHRL
EESFLVEEDKKHERHPIFGNIVDEVAYHEKYPTIYHLRKKLVDSTDKADLRLIYLALAHMIKFRGHFLIE
GDLNPDNSDVDKLFIQLVQTYNQLFEENPINASGVDAKAILSARLSKSRRLENLIAQLPGEKKNGLFGNL
IALSLGLTPNFKSNFDLAEDAKLQLSKDTYDDDLDNLLAQIGDQYADLFLAAKNLSDAILLSDILRVNTE
ITKAPLSASMIKRYDEHHQDLTLLKALVRQQLPEKYKEIFFDQSKNGYAGYIDGGASQEEFYKFIKPILE
KMDGTEELLVKLNREDLLRKQRTFDNGSIPHQIHLGELHAILRRQEDFYPFLKDNREKIEKILTFRIPYY
VGPLARGNSRFAWMTRKSEETITPWNFEEVVDKGASAQSFIERMTNFDKNLPNEKVLPKHSLLYEYFTVY
NELTKVKYVTEGMRKPAFLSGEQKKAIVDLLFKTNRKVTVKQLKEDYFKKIECFDSVEISGVEDRFNASL
GTYHDLLKIIKDKDFLDNEENEDILEDIVLTLTLFEDREMIEERLKTYAHLFDDKVMKQLKRRRYTGWGR
LSRKLINGIRDKQSGKTILDFLKSDGFANRNFMQLIHDDSLTFKEDIQKAQVSGQGDSLHEHIANLAGSP
AIKKGILQTVKVVDELVKVMGRHKPENIVIEMARENQTTQKGQKNSRERMKRIEEGIKELGSQILKEHPV
ENTQLQNEKLYLYYLQNGRDMYVDQELDINRLSDYDVDHIVPQSFLKDDSIDNKVLTRSDKNRGKSDNVP
SEEVVKKMKNYWRQLLNAKLITQRKFDNLTKAERGGLSELDKAGFIKRQLVETRQITKHVAQILDSRMNT
KYDENDKLIREVKVITLKSKLVSDFRKDFQFYKVREINNYHHAHDAYLNAVVGTALIKKYPKLESEFVYG
DYKVYDVRKMIAKSEQEIGKATAKYFFYSNIMNFFKTEITLANGEIRKRPLIETNGETGEIVWDKGRDFA
TVRKVLSMPQVNIVKKTEVQTGGFSKESILPKRNSDKLIARKKDWDPKKYGGFDSPTVAYSVLVVAKVEK
GKSKKLKSVKELLGITIMERSSFEKNPIDFLEAKGYKEVKKDLIIKLPKYSLFELENGRKRMLASAGELQ
KGNELALPSKYVNFLYLASHYEKLKGSPEDNEQKQLFVEQHKHYLDEIIEQISEFSKRVILADANLDKVL
SAYNKHRDKPIREQAENIIHLFTLTNLGAPAAFKYFDTTIDRKRYTSTKEVLDATLIHQSITGLYETRID
LSQLGGDHPQIIKKGPGIDLSQLGGDSGGSGGSGGSTNLSDIIEKETGKQLVIQESILMLPEEVEEVIGN
KPESDILVHTAYDESTDENVMLLTSDAPEYKPWALVIQDSNGENKIKMLSGGSPKKKRKVPGYPYDVPDY
AGSAAPAAKRVKLDGGSGGGSGGGSGSGPAAKRVKLDGGSGGGSGGGSGSGPAAKRVKLDGPRVIILEHH
HHHH

CE_16_eA3A

MPKKKRKVDKKYSIGLAIGTNSVGWAVITDEYKVPSKKFKVLGNTDRHSIKKNLIGALLFDSGETAEATR
LKRTARRRYTRRKNRICYLQEIFSNEMAKVDDSFFHRLEESFLVEEDKKHERHPIFGNIVDEVAYHEKYP
TIYHLRKKLVDSTDKADLRLIYLALAHMIKFRGHFLIEGDLNPDNSDVDKLFIQLVQTYNQLFEENPINA
SGVDAKAILSARLSKSRRLENLIAQLPGEKKNGLFGNLIALSLGLTPNFKSNFDLAEDAKLQLSKDTYDD
DLDNLLAQIGDQYADLFLAAKNLSDAILLSDILRVNTEITKAPLSASMIKRYDEHHQDLTLLKALVRQQL
PEKYKEIFFDQSKNGYAGYIDGGASQEEFYKFIKPILEKMDGTEELLVKLNREDLLRKQRTFDNGSIPHQ
IHLGELHAILRRQEDFYPFLKDNREKIEKILTFRIPYYVGPLARGNSRFAWMTRKSEETITPWNFEEVVD
KGASAQSFIERMTNFDKNLPNEKVLPKHSLLYEYFTVYNELTKVKYVTEGMRKPAFLSGEQKKAIVDLLF
KTNRKVTVKQLKEDYFKKIECFDSVEISGVEDRFNASLGTYHDLLKIIKDKDFLDNEENEDILEDIVLTL
TLFEDREMIEERLKTYAHLFDDKVMKQLKRRRYTGWGRLSRKLINGIRDKQSGKTILDFLKSDGFANRNF
MQLIHDDSLTFKEDIQKAQVSGQGDSLHEHIANLAGSPAIKKGILQTVKVVDELVKVMGRHKPENIVIEM
ARENQTTQKGQKNSRERMKRIEEGIKELGSQILKEHPVENTQLQNEKLYLYYLQNGRDMYVDQELDINRL
SDYDVDHIVPQSFLKDDSIDNKVLTRSDKNRGKSDNVPSEEVVKKMKNYWRQLLNAKLITQRKFDNLTKA

ERGGLSELDKAGFIKRQLVETRQITKHVAQILDSRMNTKYDENDKLIREVKVITLKSKLVSDFRKDFQFY
KVREINNYHHAHDAYLNAVVGTALIKKYPKLESEFVYGDYKVYDVRKMIAKSEQEIGKATAKYFFYSNIM
NFFKHMSGSETPGTSESATPESEASPASGPRHLMDPHIFTSNFNNGIGRHKTYLCYEVERLDNGTSVKMD
QHRGFLHGQAKNLLCGFYGRHAELRFLDLVPSLQLDPAQIYRVTWFISWSPCFSWGCAGEVRAFLQENTH
VRLRIFAARIYDYDPLYKEALQMLRDAGAQVSIMTYDEFKHCWDTFVDHQGCPFQPWDGLDEHSQALSGR
LRAILQNQGNGSGSETPGTSESATPESPWETNGETGEIVWDKGRDFATVRKVLSMPQVNIVKKTEVQTGG
FSKESILPKRNSDKLIARKKDWDPKKYGGFDSPTVAYSVLVVAKVEKGKSKKLKSVKELLGITIMERSSF
EKNPIDFLEAKGYKEVKKDLIIKLPKYSLFELENGRKRMLASAGELQKGNELALPSKYVNFLYLASHYEK
LKGSPEDNEQKQLFVEQHKHYLDEIIEQISEFSKRVILADANLDKVLSAYNKHRDKPIREQAENIIHLFT
LTNLGAPAAFKYFDTTIDRKRYTSTKEVLDATLIHQSITGLYETRIDLSQLGGDHPQIIKKGPGIDLSQL
GGDSGGSGGSGGSTNLSDIIEKETGKQLVIQESILMLPEEVEEVIGNKPESDILVHTAYDESTDENVMLL
TSDAPEYKPWALVIQDSNGENKIKMLSGGSPKKKRKVPGYPYDVPDYAGSAAPAAKRVKLDGGSGGGSGS
GPAAKRVKLDGPRLEVLFQGPGLEHHHHHH

CE_2_eA3A

MPKKKRKVDKKYSIGLAIGTNSVGWAVITDEYKVPSKKFKVLGNTDRHSIKKNLIGALLFDSGETAEATR
LKRTARRRYTRRKNRICYLQEIFSNEMAKVDDSFFHRLEESFLVEEDKKHERHPIFGNIVDEVAYHEKYP
TIYHLRKKLVDSTDKADLRLIYLALAHMIKFRGHFLIEGDLNPDNSDVDKLFIQLVQTYNQLFEENPINA
SGVDAKAILSARLSKSRRLENLIAQLPGEKKNGLFGNLIALSLGLTPNFKSNFDLAEDAKLQLSKDTYDD
DLDNLLAQIGDQYADLFLAAKNLSDAILLSDILRVNTEITKAPLSASMIKRYDEHHQDLTLLKALVRQQL
PEKYKEIFFDQSKNGYAGYIDGGASQEEFYKFIKPILEKMDGTEELLVKLNREDLLRKQRTFDNGSIPHQ
IHLGELHAILRRQEDFYPFLKDNREKIEKILTFRIPYYVGPLARGNSRFAWMTRKSEETITPWNFEEVVD
KGASAQSFIERMTNFDKNLPNEKVLPKHSLLYEYFTVYNELTKVKYVTEGMRKPAFLSGEQKKAIVDLLF
KTNRKVTVKQLKEDYFKKIECFDSVEISGVEDRFNASLGTYHDLLKIIKDKDFLDNEENEDILEDIVLTL
TLFEDREMIEERLKTYAHLFDDKVMKQLKRRRYTGWGRLSRKLINGIRDKQSGKTILDFLKSDGFANRNF
MQLIHDDSLTFKEDIQKAQVSGQGDSLHEHIANLAGSPAIKKGILQTVKVVDELVKVMGRHKPENIVIEM
ARENQTTQKGQKNSRERMKRIEEGIKELGSQILKEHPVENTQLQNEKLYLYYLQNGRDMYVDQELDINRL
SDYDVDHIVPQSFLKDDSIDNKVLTRSDKNRGKSDNVPSEEVVKKMKNYWRQLLNAKLITQRKFDNLTKA
ERGGLSELDKAGFIKRQLVETRQITKHVAQILDSRMNTKYDENDKLIREVKVITLKSKLVSDFRKDFQFY
KVREINNYHHAHDAYLNAVVGTALIKKYPKLESEFVYGDYKVYDVRKMIAKSEQEIGKATAKYFFYSNIM
NFFKHMSGEASPASGPRHLMDPHIFTSNFNNGIGRHKTYLCYEVERLDNGTSVKMDQHRGFLHGQAKNLL
CGFYGRHAELRFLDLVPSLQLDPAQIYRVTWFISWSPCFSWGCAGEVRAFLQENTHVRLRIFAARIYDYD
PLYKEALQMLRDAGAQVSIMTYDEFKHCWDTFVDHQGCPFQPWDGLDEHSQALSGRLRAILQNQGNGSPW
ETNGETGEIVWDKGRDFATVRKVLSMPQVNIVKKTEVQTGGFSKESILPKRNSDKLIARKKDWDPKKYGG
FDSPTVAYSVLVVAKVEKGKSKKLKSVKELLGITIMERSSFEKNPIDFLEAKGYKEVKKDLIIKLPKYSL
FELENGRKRMLASAGELQKGNELALPSKYVNFLYLASHYEKLKGSPEDNEQKQLFVEQHKHYLDEIIEQI
SEFSKRVILADANLDKVLSAYNKHRDKPIREQAENIIHLFTLTNLGAPAAFKYFDTTIDRKRYTSTKEVL
DATLIHQSITGLYETRIDLSQLGGDHPQIIKKGPGIDLSQLGGDSGGSGGSGGSTNLSDIIEKETGKQLV
IQESILMLPEEVEEVIGNKPESDILVHTAYDESTDENVMLLTSDAPEYKPWALVIQDSNGENKIKMLSGG
SPKKKRKVPGYPYDVPDYAGSAAPAAKRVKLDGGSGGGSGSGPAAKRVKLDGPRLEVLFQGPGLEHHHHH
H

2) sgRNA sequences

| Site | Sequence |
| --- | --- |
| *AAVS1* | GTCCCCTCCACCCCACAGTGGGG |
| *FANCF site1* | GGAATCCCTTCTGCAGCACCTGG |
| *VEGFA site2* | GACCCCCTCCACCCCGCCTCCGG |
| *MSSK1-M-c* | CGTCGCCGATCTTCACAGGGTGG |
| *EMX1 site* | GAGTCCGAGCAGAAGAAGAAGGG |
| *FANCF-M-b* | AAGTTCGCTAATCCCGGAACTGG |
| *PPP1R12C site 3* | GACCCTCAGCCAGTGCTGCTCGG |
| *RNF2* | GTCATCTTAGTCATTACCTGAGG |
| *PPP1R12C site 6* | GGGGCTCAACATCGGAAGAGGGG |
| *1620* | TTTATCACAGGCTCCAGGAAGGG |
| *1618* | TTGCTTTTATCACAGGCTCCAGG |
| *Sa_Site5* | TCTGCTTCTCCAGCCCTGGC CTGGGT |
| *Sa_Site6* | GATGTTCCAATCAGTACGCA GAGAGT |

3) Expression plasmids used in this study

| Plasmids | Comments |
| --- | --- |
| pET21a-CE_16_eA3A | CE_eA3A with 16 a.a. N / 16 a.a. C linker |
| pET21a-CE_7_16_eA3A | CE_eA3A with 7 a.a. N / 16 a.a. C linker |
| pET21a-CE_2_16_eA3A | CE_eA3A with 2 a.a. N / 16 a.a. C linker |
| pET21a-CE_2_7_eA3A | CE_eA3A with 2 a.a. N / 7 a.a. C linker |
| pET21a-CE_2_eA3A | CE_eA3A with 2 a.a. N / 2 a.a. C linker |
| pET21a-eA3A-BE3 | BE3 comprising eA3A linked to the N-terminus of nCas9 |
